# Energy transfer in ubiquitous rhodopsin pumps with xanthophyll antennas

**DOI:** 10.1101/2022.08.24.505090

**Authors:** Ariel Chazan, Ishita Das, Takayoshi Fujiwara, Shunya Murakoshi, Wataru Shihoya, Andrey Rozenberg, Ana Molina-Márquez, Shirley Larom, Alina Pushkarev, Partha Malakar, Sanford Ruhman, Masumi Hasegawa, Yuya Tsukamoto, Tomohiro Ishizuka, Masae Konno, Takashi Nagata, Keiichi Inoue, Yosuke Mizuno, Kota Katayama, Rei Abe-Yoshizumi, Hideki Kandori, Rosa M. León, Susumu Yoshizawa, Mordechai Sheves, Osamu Nureki, Oded Béjà

## Abstract

Energy transfer from light-harvesting ketocarotenoids to light-driven proton pumps xanthorhodopsins has been previously demonstrated in two unique cases: an extreme halophilic bacterium^1^ and a terrestrial cyanobacterium^2^. Attempts to find carotenoids that bind and transfer energy to rhodopsin proton pumps from the abundant marine and freshwater photoheterotrophs have thus far failed^3–5^. Here, using functional metagenomics combined with chromophore extraction from the environment, we detected light energy transfer from the widespread hydroxylated carotenoids zeaxanthin and lutein to the retinal moiety of xanthorhodopsins and proteorhodopsins. The light-harvesting carotenoids transfer up to 42% of the harvested energy in the violet/blue-light range to the green-light absorbing retinal chromophore. Our data suggest that these antennas have a significant impact on rhodopsin phototrophy in the world’s lakes, seas and oceans.

## Introduction

Proteorhodopsins (PRs) are light-driven proton pumps that were first discovered in abundant marine Gammaproteobacteria via the use of metagenomics^6,7^. Since their discovery, numerous homologous proteins have been identified in diverse bacterial and some archaeal and eukaryotic groups from marine and freshwater environments^8,9^, and it was estimated that >50% of prokaryotes in the ocean’s photic zone possess these rhodopsins^10,11^. Furthermore, it was suggested that PRs are a major light-harvesting mechanism in the surface ocean^12^.

Whereas many PRs absorb light preferentially in the green region (green-absorbing-PRs (GPRs), absorption maximum (*λ*_max_) ∼520 nm), others show blue-shifted absorption (blue-absorbing-PRs (BPRs), *λ*_max_ ∼490 nm)^7,13^. This division is determined mainly by the residue at position 105 and enables PRs to absorb light according to water depth as blue light penetrates deeper in clear oceanic waters^7,14^.

The first representative of xanthorhodopsins (XRs), a family of light-driven proton pumps related to PRs, was discovered in the genome of the extreme halophilic bacterium *Salinibacter ruber*^1^. Biophysical characterization revealed that XR uses two chromophores to absorb light: an all-*trans* retinal moiety covalently bound within the retinal binding pocket inside the protein, and a 4-ketocarotenoid antenna (salinixanthin) that binds the protein from the outside and transfers light energy directly to the retinal molecule through a fenestration^15^. This way, salinixanthin enables XR to absorb a broader range of wavelengths. Another member of the family, GR from the thylakoid-less terrestrial cyanobacterium *Gloeobacter violaceus*^2^, also recruits a 4-ketocarotenoid antenna (echinenone) to transfer light energy to the retinal chromophore within the rhodopsin^16^. The 4-keto group of salinixanthin and echinenone was shown to be crucial for both binding and energy transfer^15^.

The ability to use a carotenoid antenna requires the presence of the above mentioned fenestration in the rhodopsin apoprotein that enables the energy transfer between the rings of the two chromophores, and this in turn is facilitated by a conserved Gly residue in TM5 (Gly156 in *S. ruber*’s XR)^2^. Environmental XRs (as well as PRs) are variable at this position, with the majority of proteins having either bulky Trp or Phe residues, which render the use of a carotenoid antenna impossible, or the small Gly residue. Carotenoids are abundant and diverse in nature; nevertheless, besides the two mentioned XRs, no other native microbial rhodopsins (including other XRs) could be shown to absorb light with pre-defined carotenoid antennas^3–5^.

## Results & Discussion

Inspired by the known XR/keto-carotenoid light-harvesting systems, we designed a strategy to search for rhodopsin/carotenoid complexes originating from aquatic microorganisms. For the first exploratory series of experiments, we picked a rhodopsin with fenestration discovered using functional metagenomics^17–19^ in a freshwater lake (Lake Kinneret, Israel) and exposed it to a concentrated chromophore extract from the same lake (Fig. 1a, see METHODS). As the test protein we chose rhodopsin Kin4B8 [GenBank OP056329], a member of the XR family with a Gly at XR position 156 from an uncultured Bdellovibrionota bacterium. We incubated the purified protein with the Kinneret chromophore Extract (KE, Extended Data Fig. 1) and observed a shift in *λ*_max_ after purification of the complex (Fig. 1b), suggesting a strong binding of specific chromophores. A pronounced circular dichroism (CD) spectrum characterized by carotenoid chromophores and probably retinal bands was observed upon addition of the KE cocktail to purified Kin4B8 protein (Fig. 1c), further supporting specific binding of chromophores to the rhodopsin. HPLC-DAD analysis of the enriched complex showed that the chromophores mainly consisted of the xanthophylls lutein and zeaxanthin (Fig. 1d). This result was unexpected as both pigments have a hydroxyl group in position 3 on their β- (and also ε-in the case of lutein) rings, while only 4-ketocarotenoids were previously shown to bind to rhodopsins (compare structures in Fig. 1e).

**Fig. 1.**
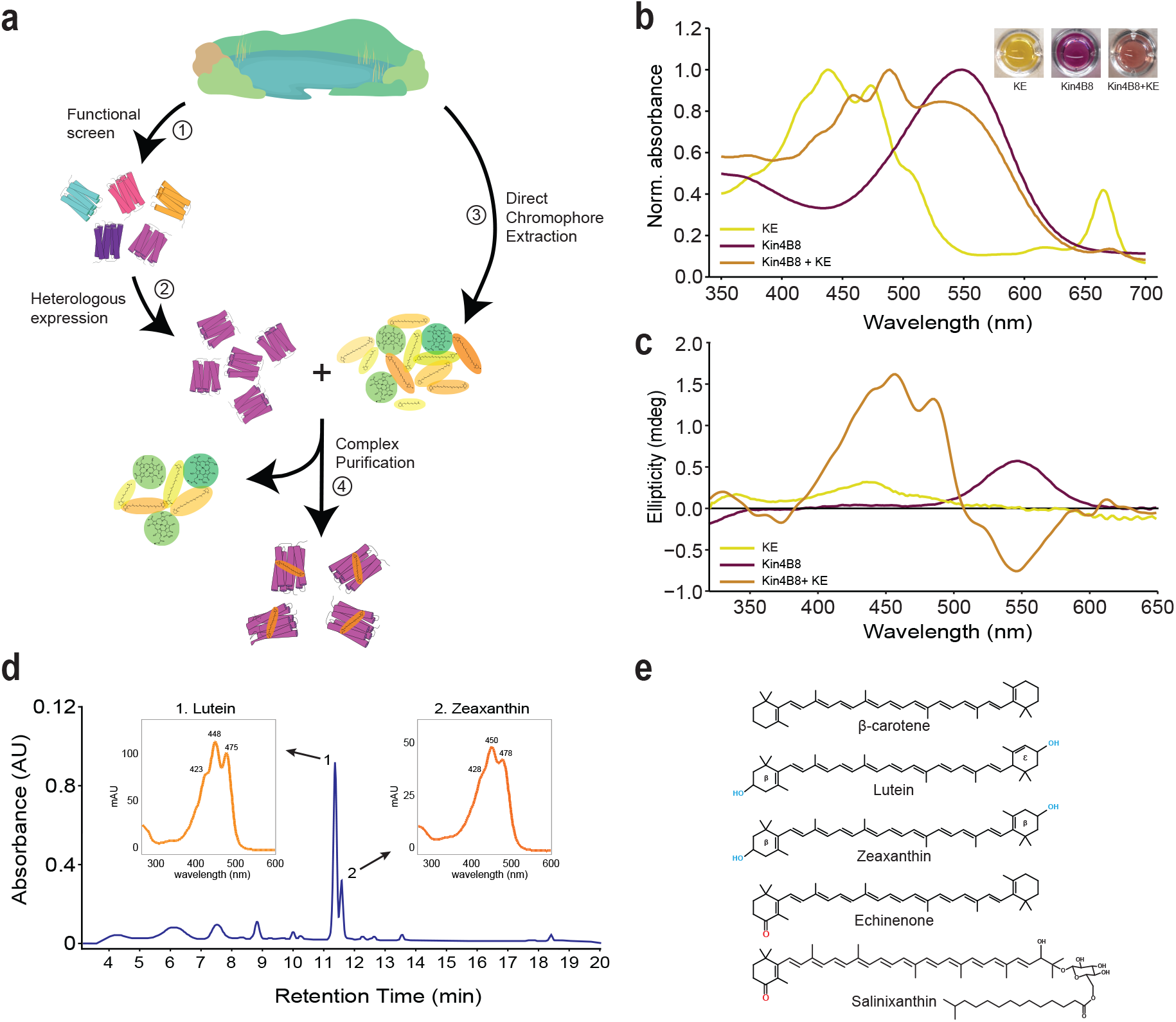
Environmental xanthophylls bind a freshwater XR. **a**, Working pipeline for searching rhodopsin/carotenoid complexes. (1) Detection of diverse rhodopsins via functional metagenomics. (2) Heterologous expression of selected rhodopsin in *Escherichia coli* and (3) incubation with chromophore extract derived from the same environment. (4) Rhodopsin is then purified and examined for changes in absorbance spectra. **b**, Absorbance spectra for the Kinneret extract (KE), rhodopsin Kin4B8, and rhodopsin Kin4B8 enriched with KE chromophores. Pictures of the samples are presented in the upper right corner. **c**, CD spectra with and without KE. **d**, HPLC-DAD chromatogram, registered at 450 nm, of the KE derived carotenoids bound to Kin4B8 protein, and UV–Vis spectra (inset) of the identified peaks, which correspond to (1) lutein and (2) zeaxanthin. **e**, 2D structures of β-carotene and various hydroxy and keto-carotenoids.

After confirming that commercially available lutein and zeaxanthin also showed binding to Kin4B8 (Fig. 2a,b, and Extended Data Fig. 2a,b), we switched to the pure chromophores for further experiments (see Fig. 1d). Fluorescence excitation spectrum (monitored at the retinal emission, 720 nm) of the Kin4B8-xanthophyll complexes exhibited three characteristic peaks of the carotenoid in addition to the expected 560 nm band of the retinal chromophore (Fig. 2c and Extended Data Fig. 2c). The occurrence of the carotenoid bands at the retinal emission indicates that excitation energy from light absorption by the carotenoid is transferred to the excited singlet state of retinal (S_1_) and contributes to emission from this state. This observation demonstrates direct energy transfer from lutein and zeaxanthin to the retinal molecule in Kin4B8. Based on the fluorescence excitation spectral measurements, we estimate that up to 40% of the light energy harvested by the carotene antenna lutein or zeaxanthin is transferred to the retinal chromophore (Extended Data Table 1). Further evidence of the energy transfer was obtained by femtosecond transient absorption (fTA) study (see METHODS for details), with the energy transfer efficiency estimated as ∼55% for the Kin4B8-lutein system (Extended Data Fig. 3). Excitation Energy Transfer (EET) from carotenoids (as assessed by fluorescence measurements with lutein) was most efficient in the 430-460 nm region (see Extended Data Fig. 4). Light-induced difference UV-visible and FTIR spectra of Kin4B8 were similar with and without lutein at 77 K (Extended Data Fig. 5), indicating that carotenoid binding does not affect retinal isomerization. Interestingly, rhodopsin Kin4B8 could also bind salinixanthin (Extended Data Fig. 6a,b), however, no energy transfer was observed in this case (Extended Data Fig. 6c). β-carotene, on the other hand, did not show binding to rhodopsin Kin4B8 (Extended Data Fig. 6d,e).

**Fig. 2.**
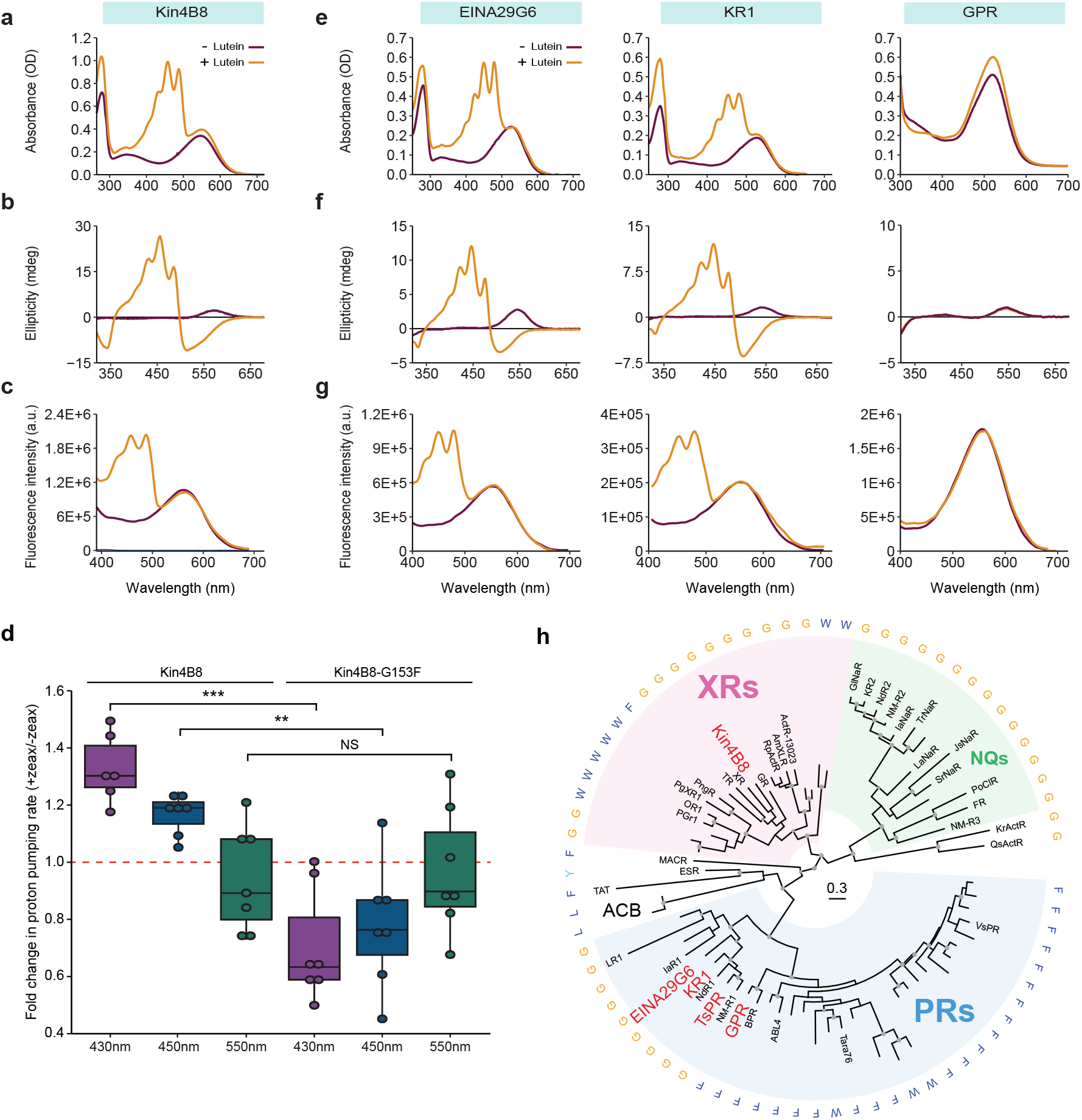
Biophysical characterization of diverse rhodopsins bound to xanthophylls. **a** and **e**, Absorbance change of different rhodopsins upon incubation with lutein. **b** and **f**, CD spectra with and without lutein. **c** and **g**, Fluorescence excitation spectra with and without lutein; emission monitored at 720 nm. See Extended Data Fig. 2 for the biophysical characterization of the same rhodopsins bound to zeaxanthin. **d**, Light-driven initial proton pumping ratios in Kin4B8 and Kin4B8-G153F *E. coli* spheroplasts with and without zeaxanthin under violet, blue and green light. Red dashed line represents proton-pumping activity when measured without zeaxanthin. Indicated are significance levels based on *t*-tests with FDR correction: *** – adjusted *p* value < 0.001, ** – adjusted *p* < 0.01. **h**, Phylogenetic tree of the clade including PRs, XRs and related families. For each one of representative sequences the residue at the fenestration position is indicated. The tree is rooted. Proteins investigated in this study are highlighted in red. Rhodopsin clade abbreviations: ACB – Archaea clade B, ESR – *Exiguobacterium sibiricum* rhodopsin, MACR – marine actinobacterial clade rhodopsins, NQs - NQ chloride and sodium rhodopsin pumps, PRs - proteorhodopsins, TAT - TAT rhodopsins, XRs - xanthorhodopsins. See Extended Data Fig. 10a for more details.

**Fig. 3.**
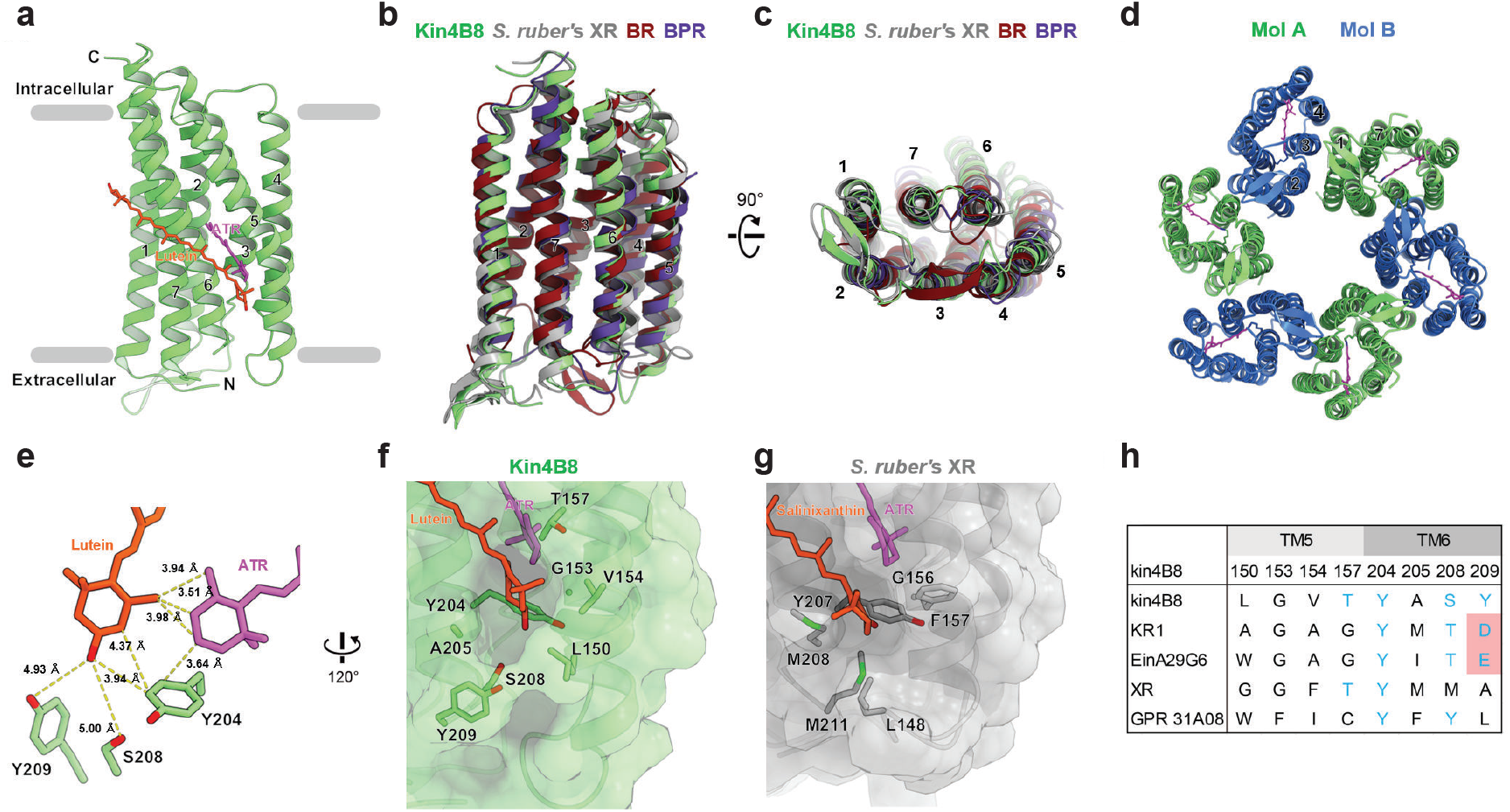
Structure of Kin4B8 XR bound to lutein. **a**, Overall structure of the monomeric unit, with lutein and retinal chromophores. **b**,**c**, Structural comparison of Kin4B8 with *S. ruber’*s XR (PDB ID: 3DDL), bacteriorhodopsin (BR) (PDB ID: 1C3W), and BPR Med12 (PDB ID: 4JQ6), with root mean square deviations (RMSD) of 1.44, 1.96, and 2.61 Å, respectively. Notably, the N-terminal region (residues 6-11) and ECL1 form a 3-stranded antiparallel β-sheet, as in *S. ruber*’s XR and other omega rhodopsins. **d**, Hexameric structure of Kin4B8, viewed from the extracellular side. Two protomers in the asymmetric unit form a hexamer with 3-fold symmetry. **e**, Positional relationship with hydroxyl ring of lutein, retinal, and surrounding residues. **f, g**, Fenestrations in Kin4B8 (**f**) and *S. ruber’*s XR (**g**). **h**, Conservation of the residues constituting the fenestrations in XRs and PRs.

**Fig. 4.**
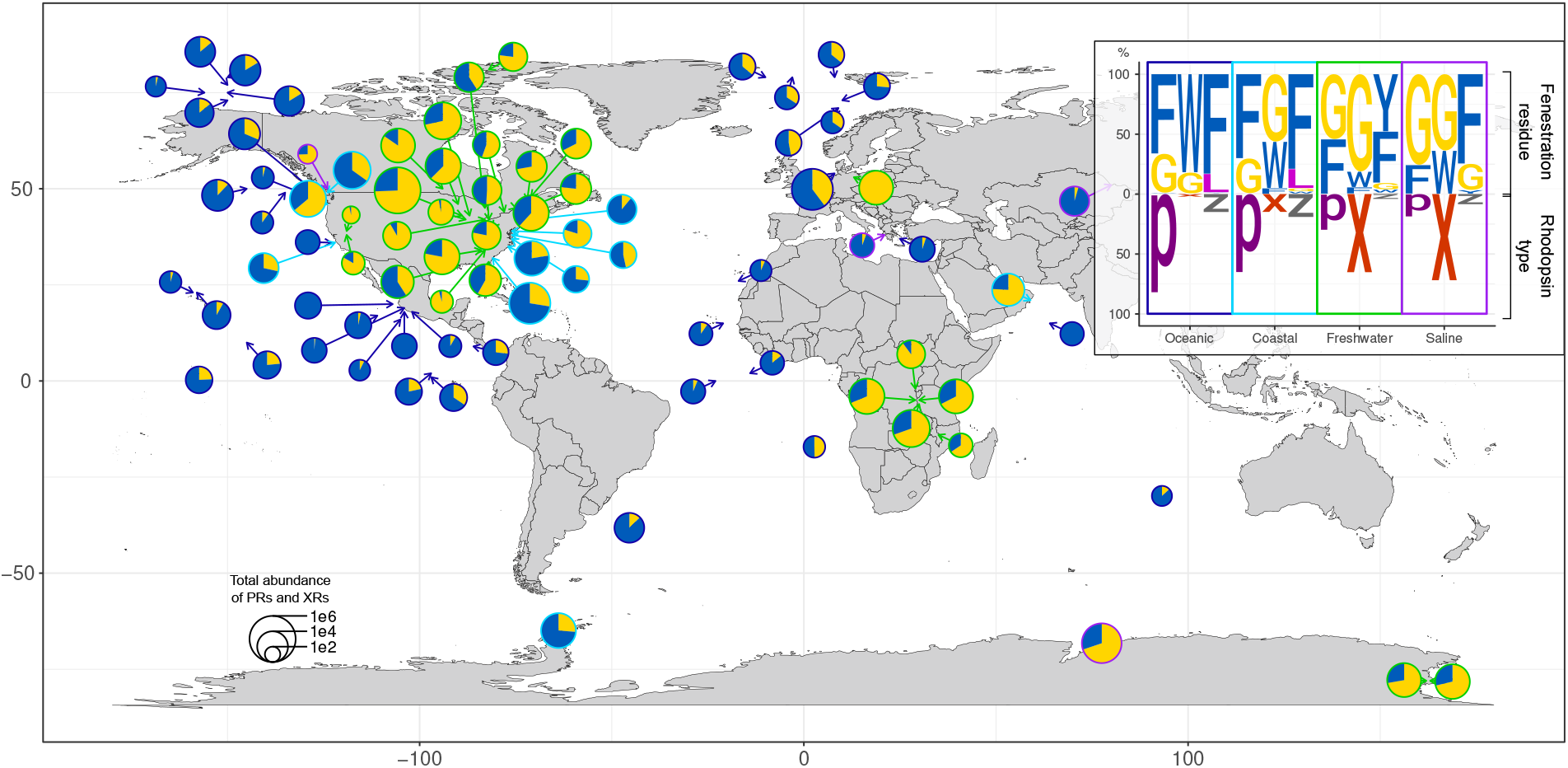
Global distribution of fenestrated XRs and PRs. Rhodopsin sequences from JGI metagenomic datasets were classified into PRs and XRs, and binned into those having bulky residues (Phe or Trp) or Gly at XR position 156. Ratios between fenestrated (yellow) and non-fenestrated (blue) PRs and XRs by location are based on the cumulative abundance for the contigs containing the corresponding rhodopsin genes and are shown as pie charts with the size of the charts proportional to the total abundance of PR- and XR-containing contigs. Locations are subdivided by habitat into: oceanic (blue stroke), coastal (cyan), freshwater (green) and inland saline (purple). Inset: ratios between PRs and XRs (x) and other superclade-P rhodopsins (z) and the incidence of the different residues at XR position Gly 156.

HPLC analysis of retinal oxime produced by the reaction between hydroxylamine and the retinal chromophore in Kin4B8 showed almost equal amounts of all-*trans* and 13-*cis* forms, independently of the light conditions, but the binding of lutein increased the fraction of the all-*trans* form (Extended Data Fig. 7a). Transient absorption changes representing red-shifted (K, O) and blue-shifted (M) photointermediates were observed by laser flash photolysis of Kin4B8 (Extended Data Fig. 7b). Multi-exponential analysis identified nine photointermediates having different absorption spectra and involving the L and N intermediates with *λ*_max_ close to the initial state (Extended Data Fig. 7c). Interestingly, a sharp peak at ∼498 nm was observed when lutein was bound to the protein (Extended Data Fig. 7b, bottom and Extended Data Fig. 7c), indicating that a large conformational change of the protein occurs during the photocycle that alters the absorption of lutein (Extended Data Fig. 7d). The transient absorption signal was enhanced for Kin4B8 bound to lutein compared with the protein without lutein at excitation wavelengths < 500 nm, indicating that energy transfer from lutein to retinal enhances retinal isomerization (Extended Data Fg. 7e).

When measured under violet (410-430 nm) or blue (440-460 nm) illumination, light-dependent outward proton flux in *E. coli* spheroplasts expressing XR Kin4B8 was enhanced upon the addition of zeaxanthin (an addition of ∼33% in violet light and ∼17% in blue light; Fig. 2d and Extended Data Fig. 8). The enhancement was not observed under green (550 nm) light illumination. This indicates that violet/blue light energy is absorbed by the xanthophyll antennas and enhances rhodopsin activity. As expected, mutating Kin4B8 Gly153 (XR position Gly156) to Phe, blocked the fenestration and abolished carotenoid binding (Extended Data Fig. 9), indicating that the fenestration in rhodopsin Kin4B8 is crucial for xanthophyll binding and energy transfer. In addition, we noticed that blocking the fenestration reduced the pumping activity significantly in the presence of zeaxanthin (Fig. 2d). This decline is likely caused by non-specific carotenoid adsorption to the spheroplasts’ surface, resulting in masking of light. This suggests that the observed increase in proton flux in wild-type Kin4B8 under violet/blue light, is in effect underestimated.

Next, we tested whether members of the PR family would be able to bind carotenoids as well. We selected the Gly156-containing PRs EinA29G6 [GenBank UJI09384] from an uncultured freshwater flavobacterium^19^ and KR1 [GenBank BAN14807] from the marine flavobacterium *Dokdonia eikasta*^20^, and GPR 31A08 [GenBank AAG10475] from a marine gammaproteobacterium^7^, which has Phe at XR position 156 instead (Fig. 2h, Extended Data Fig. 10a). Fenestrated PRs EinA29G6 and KR1 bound hydroxylated carotenoids (Fig. 2e,f, Extended Data Fig. 2d,e) and showed direct energy transfer (40% and 35%, respectively) from the xanthophylls to the retinal chromophore (Fig. 2g, Extended Data Fig. 2f and Extended Data Table 1). As expected, non-fenestrated GPR 31A08 did not bind xanthophylls, and hence did not show energy transfer.

Zeaxanthin and lutein are widespread xanthophylls in freshwater and marine environments^21–23^. Zeaxanthin is readily synthesized by a range of bacteria and is, in particular, one of the dominant carotenoids among Bacteriodota^24^, which is correlated with the abundance of fenestrated rhodopsins in this group (see Extended Data Fig. 10b). In order to demonstrate that xanthophylls can function as antenna chromophores natively, we turned to a simple cultured system. *Tenacibaculum* sp. SG-28C, a cultured marine flavobacterium, contains only a single rhodopsin (TsPR, GenBank # PQJ23084) gene in its genome (GenBank # MQVY01000000) and produces zeaxanthin as its sole carotenoid (Extended Data Fig. 11a). We demonstrate that zeaxanthin indeed binds TsPR and transfers energy to the retinal molecule (Extended Data Fig. 11b-d).

To examine xanthophylls binding to rhodopsins, we determined the crystal structure of Kin4B8 with lutein at a 3.0-Å resolution (Fig. 3a, Extended Data Fig 12a, and Extended Data Table 2). The overall structure of Kin4B8 superimposed well on that of the related XR from *S. ruber* (PDB ID: 3DDL)^25^ (Fig. 3bc). Kin4B8 has conserved structural features observed in other outward proton pumps (Extended Data Fig. 12b) and forms a hexamer in the crystal (Fig. 3d and Extended Data Fig. 12c). The oligomeric structure of Kin4B8 is very similar to that of BPR Med12 that forms a hexamer as well (Extended Data Fig. 12d)^26^, but differs from other omega rhodopsins that form pentamers instead^27^. Lutein lies transversely against the outer surface of TM6 and the angle between the two chromophore axes is about 50° (Fig. 3a). The ring moiety of lutein fits into the fenestration in Kin4B8 and forms van der Waals interactions with part of the retinal β-ionone ring (Fig. 3e, f and Extended Data Fig. 8e). The binding mode of lutein is similar to that of salinixanthin in *S. ruber*’s XR^25^ (Fig. 3g) and would enable energy transfer by a similar mechanism. The hydroxyl group of lutein is proximal to those of Ser208 and Tyr209, which constitute the bottom of the fenestration (Fig. 3e, f). Homologous residues in KR1 and EinA29G6 are also polar (Fig. 3h), whereas they are hydrophobic in *S. ruber*’s XR (Fig. 3g). Taken together, the environment at the bottom of the fenestration might distinguish between hydroxylated or ketolated carotenoids selection among various XRs and PRs.

To estimate the potential global effect of xanthophylls on rhodopsin phototrophy in aquatic environments, we calculated the proportion of Gly156-containing rhodopsins in available metagenomic datasets. As the touchstone category, we focused on PRs and XRs with the canonical motif DTE that represent the vast majority of rhodopsins from this clade in the ocean and freshwater, respectively (Fig. 4, Extended Data Fig. 10c). On average, 55% of environmental XRs and 43% of PRs contain the fenestration-enabling Gly156 (Fig. 4). Interestingly, prediction of absorption spectra revealed that PR and XR variants with Gly show a tendency to absorb in green in the region of relatively longer wavelengths (Extended Data Fig. 10d). At the same time, the fenestration site has not been previously implicated in spectral tuning in rhodopsins and blocking the fenestration in Kin4B8 (see above) had no effect on λ_max_. We thus hypothesize that the observed correlation is an indication that the more red-shifted rhodopsins tend to be fenestrated to allow a broader range of absorbed wavelengths when in complex with carotenoid antennas.

Assuming that the majority of fenestrated rhodopsins recruit secondary antennas, generalizing upon our observation of 40% energy transfer from the carotenoid antenna to the retinal molecule in Kin4B8 and taking into account the higher energy of violet (3.10 eV) and blue (2.75 eV) photons compared to green (2.38 eV), we estimate that antenna-containing rhodopsins are able to harvest an additional 46-52% energy compared to antenna-less rhodopsins. Our work shows that the binding of carotene antennas to rhodopsins is a widespread phenomenon with a significant impact on rhodopsin phototrophy.

## Supporting information

Supp Methods plus Supp Figures

## Acknowledgments

We thank J.K. Lanyi for commenting on the manuscript, M. Shalev-Benami for help in optimizing protein purification, G. Tzuri and T. Isaacson for sharing materials, and J. Anton for providing *S. ruber* for the isolation of salinixanthin standard.

## Author contributions

A.C. and A.P. conceived the project; A.C. performed environmental sampling, functional metagenomics, carotene extraction, protein biochemistry, carotene binding, and light-dependent proton pumping; A.R. performed bioinformatics; S.L. performed molecular biology; T.F., M.H., Y.T. and S.Y. performed absorption, emission, and carotene characterization of the PR-containing flavobacterial isolate; T.I., M.K., T.N. and K.I. performed laser-flash photolysis, Y.M., K.K., R.A.-Y. and H.K. performed low-temperature UV-visible and FTIR spectroscopy, M.Shunya, W.S. and O.N. performed crystallography, P.M. and S.R. performed ultrafast spectroscopy, I.D. and M.Sheves performed absorption, emission and circular dichroism spectroscopies; A.M.-M. and R.M.L. performed carotene characterization from environmental samples and from the rhodopsin-bound carotenes; O.B. coordinated the project; A.C. and O.B. wrote the paper with input from all authors.

## Data availability

All data are available in the main text or the supplementary materials. The sequence of fosmid Kin4B8 was deposited in GenBank under accession # OP056329. The structure of rhodopsin Kin4B8 bound to lutein was deposited in PDB under code 7YTB. The code used for the bioinformatic analysis is available from the github repository https://github.com/BejaLab/antenna and the data are deposited in Figshare repository doi:10.6084/m9.figshare.20502384.

