## Supplementary material for "Energy transfer in ubiquitous rhodopsin pumps with xanthophyll antennas": Supp Methods plus Supp Figures

[1]Faculty of Biology, Technion – Israel Institute of Technology, Haifa 3200003, Israel. [2]Department of Molecular Chemistry and Materials Science, Weizmann Institute of Science, Rehovot 761001, Israel. [3]Atmosphere and Ocean Research Institute, The University of Tokyo, Chiba 277-8564, Japan. [4]Department of Biological Sciences, Graduate School of Science, The University of Tokyo, Tokyo 113-0033, Japan. [5]Laboratory of Biochemistry and Molecular Biology, Faculty of Experimental Sciences, Marine International Campus of Excellence (CEIMAR), University of Huelva, Huelva 21071, Spain. [6]Institute of Chemistry, The Hebrew University of Jerusalem, Jerusalem 9190401, Israel. [7]The Institute for Solid State Physics, The University of Tokyo, Chiba 277-8581, Japan. [8]Department of Life Science and Applied Chemistry, Nagoya Institute of Technology, Showa-ku, Nagoya 466-8555, Japan. [9]OptoBioTechnology Research Center, Nagoya Institute of Technology, Showa-ku, Nagoya 466-8555, Japan

[10]Present address: Institute for Biology, Experimental Biophysics, Humboldt-Universität zu Berlin, Berlin 10115, Germany. [11]Present address: Institute for Extra-cutting-edge Science and Technology Avant-garde Research (X-star), Japan Agency for Marine-Earth Science and Technology (JAMSTEC), Kanagawa 237–0061, Japan

#### Materials & Methods

Carotenes: lutein (PHR1699) and β-carotene (PHR1239) were purchased from Sigma-Aldrich, and zeaxanthin (A132185) from AmBeed. Salinixanthin was extracted from purified cell membranes of *S. ruber* containing XR as previously described[2]. Briefly, purified XR was denatured in 2% SDS, followed by lyophilization for 20 hrs. The lyophilisate was suspended in acetone by 10 minutes vortexing followed by centrifugation at 30,000 g, at 4°C, 15 min. The resultant residual material mixture after evaporating acetone, was run in a silica column, and salinixanthin was eluted and collected with 30/70 (v/v) acetone/hexane mixed solvent after washing several times with 10/90 (v/v) acetone/hexane mixed solvent.

##### Sample preparation to study carotene interaction with rhodopsins

Lutein and zeaxanthin concentrated stock solutions were prepared in DMSO. Approximately equimolar concentration of carotenoids was added to the protein solubilized in 0.05% DDM, 0.3 M NaCl, 0.05 M phosphate buffer at pH 7.5. After incubation of the carotenoids with the proteins for one day, the complex was purified using a Ni-resin column to eliminate unbound lutein or zeaxanthin. These samples were used for further spectral studies. For the measurements performed at pH 5.5, 50 mM citrate buffer (with 300 mM NaCl and 0.05% DDM) of pH 5.5 was added to the samples and pH was adjusted by the addition of a small volume of HCl or NaOH if required further.

##### Reduction of the retinal protonated Schiff-base bond in Kin4B8-lutein complex

To the purified Kin4B8-lutein complex (in 0.1 M pH 7.5 phosphate buffer with 300 mM NaCl and 0.05% DDM), 0.15 M $NaBH_4$ were added. The reaction was carried out under continuous illumination for 20 min. The light used was filtered through a long pass cut-off filter with λ >550 nm (Schott, Germany). Then unreacted $NaBH_4$ was eliminated by dialyzing the sample against 0.1 M NaCl and 0.02% DDM. The absorption spectrum of the retinal protonated Schiff-base reduced Kin4B8 with lutein is shown in Extended Data Fig. 3.

##### Rhodopsins expression

**Kin4B8-XR –** Fosmid 4B8 was selected from our previous metagenomic screen of a fosmid library from lake Kinneret[18]. The 39 kbp insert shared high sequence similarity and synteny with MAGs from undescribed genus UBA2466 (GTDB classification: phylum Bdellovibrionota, c__FAC87, o__UBA2466, f__UBA2466, g__UBA2466). The ORF of the XR gene, in particular, had a 99.5% and 99.4% identity to assemblies GCA_002359565.1 and GCA_009926805.1, respectively. The Kin4B8 XR gene was cloned into pBAD plasmid (pBAD TOPO™ TA Expression Kit by Thermo Fisher Scientific) using forward - 5'-AAACCATGGGTTCTGCAACTACACTAACGCTG-3' and reverse - 5'-GCGTTTAAACTTAGTGATGGTGATGATGAGCGGGTAGTGAGCC-3' primers containing NcoI and MssI (PmeI) restriction sites and 6X His-tag at the C-terminus. Point mutation G153F (Gly156 in *S. ruber*'s XR), prepared in order to block the fenestration in Kin4B8, was obtained using NEB Q5 site directed protocol (https://nebasechanger.neb.com/) with primers 5'-ATTAATGTGGTTCGTCTTAAGTACCGTGCCCTTCC-3' and 5'-CGAGCGCCAGCTTGGCTT-3'.

DH10b *E. coli* cells harboring pBAD-Kin4B8 or pBAD-Kin4B8-G153F plasmids, were grown in LB supplemented with 50 µg/ml ampicillin at 220 RPM at 37 °C. When the $OD_{600}$ reached 0.6, expression was induced using a 0.1 % final concentration of L-arabinose (Sigma-Aldrich, A3256). The induced culture was grown at 220 RPM overnight (>16 h) at 30°C. Then, the culture was incubated with 20 µM all-*trans* retinal for >4h in the dark.

**KR1 –** Plasmid and expression protocol was adapted from[20]. Briefly, BL21 *E. coli* cells harboring pET-21a - KR1 plasmid, were grown on LB with 50 µg/ml ampicillin at 220 RPM at 37 °C. When the $OD_{600}$ reached 0.5, expression was induced using a 0.5 mM final concentration IPTG (Inalco Pharmaceuticals, 1758-1400), and all-*trans* retinal at a final concentration of 20 µM was supplemented. The induced culture was grown at 220 RPM overnight (>16 h) at 30°C.

**EINA29G6-PR and GPR –** Fosmid EINA29G6 (Genbank accession MW650840.1) was isolated from a freshwater sample and the proton pumping activity for the encoded PR was assessed by us previously[19]. The 36 kbp fosmid is syntenic with scaffolds from freshwater flavobacterial MAGs, in particular to assembly GCA_016868755.1 (GTDB taxonomy: phylum Bacteroidota; c__Bacteroidia; o__Flavobacteriales;

f__Schleiferiaceae; g__TMED14) with which it shows 92.6% DNA identity in the PR ORF. Single DH10b *E. coli* colonies harboring pBAD-EINA29G6-PR[19] or pBAD-GPR plasmids[7], were used to inoculate LB media supplemented with 100 µg/ml ampicillin and 0.001 or 0.2% (EINA29G6 and GPR, respectively) final concentration of L-arabinose. The culture was incubated in a deep 96-well overnight (>16h) at 750 RPM at 30 °C. Then, the pooled plate content was incubated with 20 µM all-*trans* retinal for >4 h in the dark.

**TsPR –** Single C41 (DE3) *E. coli* colonies harboring pET21a-TsPR (TsPR: Genbank accession PQJ23084.1)[28] were used to inoculate LB media supplemented with 100 µg/ml ampicillin. Before protein expression, transformants were cultured at 37°C in 2× YT medium (NaCl 5 g/L, Bacto Tryptone 16 g/L, and Bacto Yeast Extract 10 g/L, pH 7.0) with 100 µg/ml ampicillin until the absorbance at 660 nm reached 0.5. Protein expressions were then induced at 37°C for 4 h in the dark by adding 0.1 mM isopropyl β-D-1-thiogalactopyranoside (IPTG; Sigma-Aldrich, USA) and 10 µM all-*trans*-retinal.

#### Protein purification

Rhodopsin expressing *E. coli* cultures were centrifuged at 5,000 g for 15 min at 4 °C and the pellet was resuspended in a buffer containing 50 mM Tris HCl pH 8.0, 5 mM $MgCl_2$, 0.1 mM PMSF (Sigma-Aldrich, P7626). The sample was disrupted by using a microfluidizer 10 passes at 60 psi. Then, the sample was centrifuged at 5,000 g for 15 min at 4 °C to pellet undisrupted cells or large cell debris. Membranes were collected by centrifuging the sample at 37,000 g for 1 h at 4 °C and resuspended in a buffer containing 50 mM Tris HCl pH 8.0, 300 mM NaCl, 5 mM imidazole, 5 mM $MgCl_2$, 10% glycerol, and 2% DDM final concentration. The sample was incubated for > 2 h at 4 °C with gentle rotation and second centrifugation at 37,000 g for 1 h at 4 °C was performed. The supernatant was then incubated with Ni-Beads (Cube Biotech, 31103) for 1 h. Beads were washed on a gravity column using a buffer containing 50 mM Tris-HCl pH 8.0, 300 mM NaCl, 10% glycerol, 0.05% DDM, and 20-50 mM imidazole. Protein was eluted from the column using a buffer containing 50 mM Tris-HCl pH 8.0, 300 mM NaCl, 10% glycerol, 0.05% DDM, and 250 mM imidazole. Eluted protein was washed using Amicon 100 KDa cutoff (UFC910008 millipore) with storage buffer 50 mM Tris-HCl pH 8.0, 300 mM NaCl, 10% glycerol, and 0.05% DDM. The protein was then flashed-freezed in liquid $N_2$ and stored at -80 °C.

TsPR-expressing *E. coli* cells were resuspended in 7 mL of buffer containing 50 mM Tris-HCl pH 8.0, 500 mM NaCl. The cells were disrupted by sonication (Branson SFX 250 Digital Sonifier, Branson Ultrasonics, USA) on ice-cold water for 5 min. Crude membranes were obtained by ultracentrifugation at 4°C (106,800 g for 30 min; Optima XPN-90 Ultracentrifuge with a SW 32Ti rotor, Beckman Coulter, USA) and solubilized with 1% DDM. The solubilized TsPR was collected by ultracentrifugation at 4°C (106,800×g for 30 min) and purified by a HisTrap FF Ni2+-NTA affinity chromatography column (GE Healthcare, England) at room temperature (~25 °C). The purified sample was concentrated, and its buffer was replaced with a new buffer containing 25 mM MOPS pH 7.2, 500 mM NaCl, 0.1% DDM using an Amicon Ultra Filter 30 kDa cutoff (Millipore) by centrifugation at 4 °C (5000×g for 20 min; MX-305, TOMY SEIKO Co., Ltd., Japan).

#### Environmental chromophores sampling and extraction

200 L of water were sampled on 24 August at 10 AM in Lake Kinneret, Station A (32° 49.27792 N, 35° 35.34253 E). Water samples were then concentrated by a Tangential Flow Filtration system (TFF, Repligen, N06-E100-05-S) with a 100 KDa cutoff after pre-filtration through a mesh net. Concentrated water samples were disrupted using a microfluidizer (Microfluidics Corporation, M-110S) by applying 30-80 PSI for 5 cycles. Samples were then freeze-dried using a lyophilizer (SCANVAC, Coolsafa 110-4) for ~48 h. Chromophore extraction was done directly on the dried material using hexane extraction[29]. Briefly, dried samples were resuspended in 10 mL acetone by applying extensive pipetting and vortexing. Hexane and 10% NaCl were added to the mixture in a 2:2:1 ratio (Acetone: Hexane: 10% NaCl). The mixture was vortexed and then centrifuged at 3,000 g at 4°C for 3 min. The hexane (top) layer was then transferred to a separate falcon and the process was repeated till the hexane phase became colorless. Combined hexane fractions were then dried using $N_2$ gas and reconstituted in 5 mL Abs. ethanol.

#### *Tenacibaculum* sp. SG-28 chromophores extraction

*Tenacibaculum* sp. SG-28 cells were cultured in 500 mL- 1/2 strength ZoBell's 2216E medium under light condition at 25 °C. Cultured cells were collected by centrifugation

(4,400 g for 10 min at 20 °C) and then washed twice in 100 mM NaCl. Intracellular pigments were extracted with acetone/methanol (7:3 v/v). Air-dried pigments were resuspended in 30 mL buffer containing 25 mM MOPS pH 7.2, 500 mM NaCl, 0.1% DDM and filtered through 0.22 µm pore size filter (Advantech, Japan) to remove insoluble pigments. The filtered solution was concentrated using an Amicon Ultra Filter 30 kDa cutoff (Millipore) by centrifugation at 15 °C (4,000×g for 60 min; MX-305, TOMY SEIKO Co., Ltd., Japan).

**Binding of chromophores extract to rhodopsin proteins**

Kin4B8 + Kinneret chromophore extract (KE) - 10 mg of Kin4B8 purified protein were mixed with 5 ml KE at OD ~5 and incubated overnight with gentle rotation at 4°C. ~2.5 ml of Ni-Beads were added to the mixture for 4 h at 4°C. The protein was then washed extensively using a storage buffer and then eluted using the same buffers used for the initial purification.

TsPR and *Tenacibaculum* sp. SG-28 intracellular pigments extract (TE) - Purified TsPR and TE were mixed in a ratio of 450 µl (A520 of 0.5) to 600 µl (A450 of 1.5) and incubated overnight with gentle rotation at 4 °C. The mixed samples were purified using by a HisTrap FF Ni2+-NTA affinity chromatography column (Takara Bio, USA) at 4 °C. The concentration and replacement of the buffer was carried out in the same way as purification of TsPR.

**Chromatographic analysis of carotenoids**

Lyophilized KE samples (10 mg) were resuspended in 1 mL of methanol, transferred into 2-mL screw-top polystyrene tubes with 0.5 g of 0.5 mm glass beads under $N_2$ atmosphere, treated for 2 min in a Genie disruptor and incubated overnight at -20°C. After centrifugation at 9,700 g for 10 min, the supernatant was analyzed in a Hitachi Chromaster HPLC equipped with a DAD detector, using an RP-18 column and a flow rate of 1 mL min$^{-1}$. Solvents were (A) a mixture of acetonitrile/water (9:1 v/v) and (B) pure ethyl acetate. The gradient elution program was as follows: 0–16 min 0%– 60% A; 16–30 min 60% A; 30–35 min 100% A. The column temperature was kept at 25°C, and the injection volume was 100 µL. Chromaster Hitachi control software was used for data processing. Salinixanthin standard was prepared by purification from *S. ruber*.

Other standards were purchased from Sigma-Aldrich (Munich, Germany) or DHI Lab (Hørsholm, Denmark).

To identify pigments bound to TsPR, pigments analyses of the TE and the TE/TsPR mixture were performed by HPLC, respectively. One hundred µl of TE (A450 of 1.0) or TE/TsPR mixture (A520 of 0.5) was added to 500 µL of methanol and evaporated at 30 °C. Air-dried samples were resuspended in 500 µL of acetone/methanol (1:9 v/v) and stored at -20 °C for 3 h, respectively. These solutions were analyzed in a Shimadzu HPLC (model Prominence-i LC-2030) equipped with a PDA detector, using a Kinetex 5 µm EVO C18 100A column (Phenomenex, USA) and a flow rate of 2 mL $min^{-1}$. Solvent was a mixture of methanol/water (9:1 v/v). The column temperature was kept at 60 °C, and the injection volume was 20 µL. LabSolutions software was used for data processing. Purchased zeaxanthin (FUJIFILM Wako Pure Chemical corporation, Japan) was used as a standard.

**Absorption spectroscopic measurements**

UV−vis absorption spectral measurements of rhodopsins (Kin4B8, Kin4B8-G153F, KR1, EINA29G6, GPR) with the carotenoids (lutein, zeaxanthin, beta-carotene, Salinixanthin) and KE were taken with a Cary 8454 UV−vis spectrophotometer (Agilent Technologies, CA). Protein and carotenoid concentration was kept in ~2-6 µM range for absorption spectral measurements.

UV−vis absorption spectral measurements of TsPR and TE/TsPR mixture were taken with a UV-2600 spectrophotometer (Shimadzu, Japan).

**Circular Dichroism (CD) spectroscopic measurements**

CD spectroscopic measurements of rhodopsins (Kin4B8, Kin4B8-G153F, KR1, EINA29G6, GPR) with the carotenoids (lutein, zeaxanthin, beta-carotene, Salinixanthin) and KE were performed with Chirascan CD spectrometer (Applied Photophysics). A quartz cell of path length 1 cm was used for the measurements. CD spectra were recorded with 2.1 nm bandwidth resolution in 1 nm interval.

CD spectroscopic measurements of TsPR and TE/TsPR were performed with a J-725 spectrometer (Jasco, Japan) at room temperature (~25 °C). A quartz cell of path length 1 cm was used for the measurements. CD spectra were recorded with 1 nm bandwidth resolution in 1 nm interval.

**Low-temperature UV-visible and FTIR spectroscopic analysis**

Low-temperature UV-visible and FTIR spectroscopies were performed as described previously[30], except for minor modifications of the reconstitution into the membrane process. Briefly, purified Kin4B8 samples, with and without lutein, were reconstituted into a membrane composed of POPE:POPG (3:1 mol/mol) (POPE:POPG; Avanti) with a protein-to-lipid molar ratio of 1:20, by removing the DDM with Bio-Beads (SM-2, Bio-Rad). Kin4B8 samples in POPE:POPG liposomes were washed repeatedly with a buffer containing 5 mM NaCl and 2 mM $NaH_2PO_4$ (pH 7.25) and collected by ultracentrifugation for 20 min at 222,000 x g at 4 °C. The lipid-reconstituted Kin4B8 sample was finally suspended in a same buffer and then placed on a $BaF_2$ window to prepare dry-layer thin film. Kin4B8 films were hydrated with 1 μL $H_2O$ before measurements. The sample was then placed in the cell of a cryostat (Optistat, Oxford Instruments) mounted in the UV-vis spectrometer (V-750, JASCO) and a FTIR spectrometer (Carry670, Agilent Technologies) and cooled to 77 K. Each difference spectrum was calculated from two spectra constructed from 128 interferograms with 2 $cm^{-1}$ resolution. Illumination with 540 nm light for 30 sec through an interference filter (KL-54 interference filter, Toshiba) at 77 K converted Kin4B8 into the K intermediate, which reverted to the original state upon illumination at >590 nm light (R-61 cut-off filter, Toshiba) for 30 sec, as evidenced by a mirror image of the difference spectra. Averages of 100 and 60 experiments were conducted for the spectra of Kin4B8 with and without lutein, respectively.

**High performance liquid chromatography (HPLC) analysis of retinal isomers**

The HPLC analysis of retinal isomers was performed as described in[31] with minor modification. Purified samples in 50 mM Tris–HCl (pH 8.0), 150 mM NaCl, 0.1% DDM, 10% Glycerol. were kept at room temperature in the dark before the experiments. For light adaptation, samples were illuminated with green light (530 ± 5 nm) for 60 s, followed by incubation in the dark for 60 s. A 75-µL sample was mixed with 280 µL of 90% (v/v) methanol aqueous solution and 25 µL of 2 M hydroxylamine ($NH_2OH$) to convert retinal chromophore into retinal oxime, and then the retinal oxime was extracted with 800 µL of *n*-hexane. A 200 µL of the extract was injected into an HPLC system equipped with a silica column (particle size 3 µm, 150 × 6.0 mm; Pack SIL, YMC, Japan), a pump (PU-4580, JASCO, Japan), and a UV–visible detector (UV-4570, JASCO, Japan). The solvent for the mobile phase was *n*-hexane containing 15 % ethyl acetate and 0.15 % ethanol and the flow rate was 1.0 mL $min^{-1}$. The molar composition of the retinal isomers in the sample was determined with the molar extinction coefficient at 360 nm for each isomer (all-*trans*-15-*syn*: 54,900 $M^{-1}$ $cm^{-1}$; all-*trans*-15-*anti*: 51,600 $M^{-1}$ $cm^{-1}$; 13-*cis*-15-*syn*, 49,000 $M^{-1}$ $cm^{-1}$; 13-*cis*-15-*anti*: 52,100 $M^{-1}$ $cm^{-1}$; 11-*cis*-15-*syn*: 35,000 $M^{-1}$ $cm^{-1}$; 11-*cis*-15-*anti*: 29,600 $M^{-1}$ $cm^{-1}$).

#### Laser flash photolysis

For the laser flash photolysis spectroscopy, Kin4B8 with and without lutein was solubilized in 50 mM Tris–HCl (pH 8.0), 150 mM NaCl, 0.1% DDM, 10% Glycerol. Optical density of the rhodopsin was adjusted to ~0.4–0.5 (protein concentration ~0.2–0.25 mg/mL) at the absorption maximum wavelengths. The laser flash photolysis measurement was conducted as previously described. The nano-second second harmonics of Nd–YAG laser ($\lambda$ = 532 nm, INDI40, Spectra-Physics, CA) was used for the excitation of Kin4B8. The transient absorption spectra were obtained by monitoring the intensity change of white-light from a Xe-arc lamp (L9289-01, Hamamatsu Photonics, Japan) passed through the sample with an ICCD linear array detector (C8808-01, Hamamatsu, Japan). To increase the signal-to-noise (S/N) ratio, 60–100 spectra were averaged, and the singular-value-decomposition (SVD) analysis was applied. To measure the time-evolution of transient absorption change at specific wavelengths, the output of a Xe-arc lamp (L9289-01, Hamamatsu Photonics, Japan) was monochromated by monochromators (S-10, SOMA OPTICS, Japan) and the change in the intensity after the photo-excitation was monitored with a photomultiplier tube (R10699, Hamamatsu Photonics, Japan). To increase S/N ratio, 100–200 signals were averaged.

To measure the transient absorption change of pyranine due to proton release and uptake by Kin4B8 wildtype, the protein was solubilized in 50 mM sodium-phosphate, 300 mM NaCl, 0.05% DDM at pH 7.5. Nano-second pulses from an optical parametric oscillator (basiScan, Spectra-Physics, CA) pumped by the third harmonics of Nd–YAG laser ($\lambda$ = 355 nm, INDI40, Spectra-Physics, CA) were used for the excitation of Kin4B8 at different wavelengths ($\lambda_{exc}$ = 432, 458, 473, 488, 547, and 595 nm). The pulse energy was lowered to 0.3 mJ/cm$^2$ to keep the linearity between the number of the absorbed photon and the transient absorption change.

**Fluorescence spectroscopic measurements**

Fluorescence emission and excitation spectral measurements were taken on a Jobin Yvon-Spex Fluorolog-3 spectrofluorometer, which is composed of a 450W Xe-lamp as light source, double-grating monochromator in the excitation and emission positions, a photomultiplier tube detector (R928P). The Slit-width of both the emission and excitation channel were mostly kept at 8 or 10 nm. Absorbance of the samples used for fluorescence measurements were kept within 0.3 OD with respect to the carotenoid absorption maximum. However, collected spectral profiles were further corrected for the internal absorption effect. Retinal fluorescence emission intensity of Kin4B8 was good enough for measurements at pH 7.5, but for KR1, GPR, and EINA29G6 it was too weak to be detectable at this pH. Therefore, for these later three rhodopsins, all the fluorescence measurements were done at pH 5.5 (below the p$K_a$ of the retinal PSB counter-ions) in order to get detectable fluorescence signals. Excitation spectra were sampled at 720/730 nm to avoid the strong Raman bands that mask the retinal fluorescence. Fluorescence excitation spectra were scaled to the respective retinal absorption spectral for the calculation of excitation energy transfer efficiency of the respective donor-acceptor pairs. A representative figure (for Kin4B8-lutein system) showing the absorption and fluorescence excitation spectral profiles scaled together is presented in Extended Data Fig. 13. The following equation, relating the fluorescence excitation spectral profile and corresponding absorption spectral profile, was used for estimation of quantum efficiency of excitation energy transfer from the carotenoid antenna to the retinal chromophore: $Exc(\lambda) = (1 - 10^{-A}) \times (A_r + \phi A_c) / A$[32], where $Exc(\lambda)$, and $A$ (=$A_r$+$A_c$) are the excitation spectrum and absorbance of the complex, respectively, $A_c$ and $A_r$ are the absorption spectrum of bound carotenoid, and the retinal component, respectively, and ϕ is the quantum efficiency of energy transfer.

Fluorescence emission and excitation spectra of TsPR and TE/TsPR were measured on the RF-6000 spectrofluorometer (Shimadzu, Japan) at room temperature (~25 °C). The Slit-width of both the emission and excitation channel were 5 nm. Scattered excitation light was blocked by a long-pass filter (>510 nm, Asahi Spectra, Japan). Absorbance of the samples used for fluorescence measurements were TsPR sample Abs525 of 0.5, and TE-TsPR sample Abs485 of 0.6, respectively. The fluorescence measurements were done at pH 5.5 and excitation spectra were sampled at 720 nm. The following equation relating the fluorescence excitation spectral profile and corresponding absorption spectral profile, was used to correct the inner filter effect (IFE): $F_{ideal} = F_{obs} 10^{Abs_{Ex}/2}$, where $F_{ideal}$ is the ideal fluorescence-signal spectrum absence of IFE, $F_{obs}$ and $Abs_{Ex}$ are the excitation and absorbance spectra, respectively[33].

#### <u>Femtosecond transient absorption measurements of Kin4B8-Lutein</u>

fTA measurements were performed using a hybrid Ti-Saphire laser system operating at 1000 Hz. A description of the laser system, pump and probe pulse generation and its detection are detailed elsewhere[34]. fTA measurements were performed on an optical cell with a 0.5 mm quartz window and 0.4 mm path length. The sample cell was rotated rapidly to prevent photodegradation. In addition, sample integrity was monitored by measuring its absorbance throughout the measurement.

Treatment of the native rhodopsin with $NaBH_4$ yielded a reduced retinal protonated Schiff-base bond (RPSB), which in turn abolishes energy transfer from the carotenoid, yet maintaining a similar retinal binding site[35,36]. In the case of the Kin4B8(reduced RPSB bond)-lutein complex, the $S_2$ state's lifetime of lutein was 140 fs ($\tau_D$), which was shortened to 60 fs ($\tau_{DA}$) for the Kin4B8-lutein complex, confirming the energy transfer process. The efficiency of energy transfer ($\phi_{ET}$) is calculated by using $\phi_{ET} = 1 - \tau_{DA}/\tau_D$. Refer to Extended Data Fig. 3 for the femtosecond decay kinetics of $S_2$ state of lutein in the complex.

#### Proton pumping measurements

The protocol was adapted from[13] with minor modifications. Briefly, 200 ml of rhodopsin-expressing *E. coli* cells were centrifuged at 3600 g for 10 min and resuspended into 20 ml of 30 mM Tris-HCl pH 8.0, and 20% sucrose. 200 µg of lysozyme (Sigma-Aldrich, L6876) was added to the cell suspension and gently rotated for 1 h at room temperature. Spheroplasts were centrifuged at 3,600 g for 15 min at room temperature and the pellet was resuspended with 2 ml of 100 mM KPi pH 7.0, 20 mM $MgSO_4$, and 20% sucrose supplemented with 4 mg of DNase I (Sigma-Aldrich, DN25). The solution was injected using a syringe into 50 ml of 50 mM KPi pH 7.0 for 15 min at 37 °C. Na-EDTA was added to the mixture to a final concentration of 10 mM and the solution was stirred for another 15 min. Then, $MgSO_4$ was added at a final concentration of 15 mM, followed by another 15 min of stirring. Cell debris were removed by centrifugation at 3,600 g for 10 min and vesicles were collected by centrifugation at 16,000 g for 1 h. Spheroplasts were resuspended with 2 ml of 100 mM KPi pH 7.0, and 10 mM $MgSO_4$. Addition of 20 µM zeaxanthin to ~ 1 ml (1/2 of the volume) followed by overnight incubation at 4°C with gentle rotation. Spheroplasts then were washed twice with 100 mM KPi pH 7.0, and 10 mM $MgSO_4$ and three times with unbuffered solution (10 mM NaCl, 10 mM $MgSO_4 \cdot 7H_2O$, 100 µM $CaCl_2$). Samples were kept in the dark until pH was stabilized and illuminated using Leica 1177 equipped with a 150W halogen lamp and filtered with violet, blue and green interference filters (420-430, 445-455, and 560-600 nm, respectively); light intensity was ~225, 335 and 990 µmol $m^{-2}$ $sec^{-1}$, measured after the interference violet, blue and green filters, respectively (measured using LI-COR Biosciences LI-250A light meter). pH was monitored using LAQUA F-72G pH/ION meter (HORIBA scientific) equipped with a 9618S-10D pH microelectrode. pH results were converted into proton concentration $[H^+] = 10^{-ph}$, and the fold change in proton pumping rate over the first ten seconds of illumination was calculated using the following equation:

$$Fold\ Change = \frac{([H^+]_{t_{10}} - [H^+]_{t_0})_{+zeaxanthin}}{([H^+]_{t_{10}} - [H^+]_{t_0})_{-zeaxanthin}}$$

Where the fold change in proton pumping rate is represented by the delta in proton concentration over the first ten seconds of illumination of the result with zeaxanthin divided by the results without zeaxanthin[37]. Differences in pumping activity between the wildtype variant and the fenestration mutant were assessed with tailed *t*-test after exploring normality with the Shapiro-Wilk test. The *p*-values were FDR adjusted.

**Protein expression and purification for structural analysis**

pBAD-Kin4B8 was transfected in *E. coli* C41 (Rosetta). The transformant was grown in LB supplemented with 50 μg/ml ampicillin and 10 μg/ml at 220 RPM at 37 °C. When the OD600 reached 0.6, expression was induced using a 0.1% final concentration of L-arabinose (Sigma-Aldrich, A3256). The induced culture was grown at 120 RPM overnight (>16 h) at 25°C. Then, the pooled plate content was incubated with 20 μM all-*trans* retinal for >4 h in the dark. The collected cells were disrupted by sonication, in buffer containing 20 mM Tris-HCl, pH 8.0, 150 mM NaCl, and 10% glycerol. The crude membrane fraction was collected by ultracentrifugation at 180,000 g for 1 h. The membrane fraction was solubilized in buffer, containing 20 mM Tris-HCl, pH 8.0, 150 mM NaCl, 1% DDM, 10% glycerol, and 10 mM imidazole, for 2 h at 4 °C. The supernatant was separated from the insoluble material by ultracentrifugation at 180,000 g for 20 min, and incubated with Ni-NTA resin (Qiagen) for 30 min. The resin was washed with ten column volumes of wash buffer, containing 20 mM Tris-HCl, pH 8.0, 500 mM NaCl, 0.03% DDM, 10% Glycerol, and 25 mM imidazole. The resin was incubated with the one column volume of wash buffer containing 100 μM lutein. Then, the resin was washed with five column volumes of wash buffer. The protein was eluted in buffer, containing 20 mM Tris-HCl, pH 8.0, 150 mM NaCl, 0.03% DDM, 10% Glycerol, and 300 mM imidazole. The eluate was dialyzed against buffer (20 mM Tris-HCl, pH 8.0, 150 mM NaCl, 0.03% DDM). The protein was concentrated to 40 mg $ml^{-1}$ using a centrifugal filter device (Amicon 50 kDa MW cut-off), and frozen until crystallization.

**Crystallization**

The protein was reconstituted into monoolein at a weight ratio of 1:1.5 (protein:lipid). The protein-laden mesophase was dispensed into 96-well glass plates in 30-nl drops and overlaid with 800 nl precipitant solution, using a Gryphon robot (ARI), as described previously[38]. Crystals of Kin4B8 were grown at 20 °C in precipitant conditions containing 25% PEG550MME, 100 mM HEPES-NaOH, pH 7.0, and 180 mM potassium thiocyanate. The crystals were harvested directly from the LCP using micromeshes (MiTeGen) and frozen in liquid nitrogen, without adding any extra cryoprotectant.

#### Data collection and structure determination

X-ray diffraction data were collected at the SPring-8 beamline BL32XU with an EIGER X 9M detector (Dectris), using a wavelength of 1.0 Å. In total, 282 small-wedge (10° per crystal) datasets using a 15 × 10-μm2 beam. The collected images were processed using KAMO[39] with XDS[40], and 45 datasets were indexed with the consistent unit cell parameters. After correlation coefficient-based clustering using normalized structure factors followed by merging using XSCALE[41] with outlier rejections implemented in KAMO, the second largest cluster consisting of 106 datasets was selected for the downstream analyses, because it gave the highest inner-shell and outer-shell CC1/2. Kin4B8 structure was determined by molecular replacement with PHASER[41], using the model calculated by AlphaFold2[42]. Subsequently, the model was rebuilt and refined using COOT[43] and phenix.refine[44]. The asymmetric unit contains two protomers (molA and molB). Lutein was modeled in molA, but could not be modeled in molB due to the discontinuous electron density. The β-ring of lutein was modeled to fit the fenestration around the retinal. The final model of Kin4B8 contained residues 6-213 and 216-259 (molA), 3-259 (molB), one lutein, 14 monoolein molecules, and 25 water molecules. Figures were prepared using CueMol (http://www.cuemol.org/ja/).

#### Bioinformatic analyses

**Rhodopsin sequence analysis.** Rhodopsin sequences were classified using a curated set of reference sequences from the superclade uniting PRs, XRs, NQ rhododopsins and related clades (PR-XR-NQ superclade). The reference sequences were used to build a custom HMM profile (the PR-XR-NQ profile)used to collect rhodopsin sequences from environmental datasets and to create a database against which query sequences were searched with usearch_global from usearch v. 11.0.667[45] for fine-grained classification. The residues at active positions were determined based on the alignment against the PR-XR-NQ profile. Absorption maxima were predicted for the query sequences using the BLASSO machine learning model published previously[46] with a custom wrapper workflow available at https://github.com/BejaLab/BLASSO-Rhodopsin.

**Metagenomic datasets.** Three metagenomic datasets were recruited to quantify the distribution of fenestrated and non-fenestrated rhodopsins: Ocean Microbial Reference Catalog (OM-RGC) v.2[47], metagenomes distributed via JGI Integrated Microbial Genomes & Microbiomes (IMG/M)[48] and Genomes from Earth's Microbiomes (GEM) catalog of metagenomically assembled genomes (MAGs)[49]. Abundance values for individual rhodopsin sequences were taken as the per-gene coverage as supplied in OM-RGC v.2 and as per-contig coverage as supplied in IMG/M. For IMG/M, only studies with unrestricted data utilization status were recruited. Individual metagenomic studies were binned into four habitat types: open ocean, coastal marine (estuaries, lagoons, channels and near-shore habitats), freshwater and inland saline water bodies. For the map visualization of the ratios between fenestrated and non-fenestrated PRs and XRs, abundances were summed for individual samples per location with a resolution of one degree.

**Phylogenetic analysis.** Phylogenetic tree was constructed for a reference set of sequences composed mostly of characterized sequences by aligning them with hmmalign from the hmmer v. 3.3.2[50] package against the reference PR-XR-NQ profile, removing unaligned residues and building the phylogeny with iqtree2 v. 2.2.0.3 with ultrafast bootstrap[51,52].

### Supplementary Information

**a**

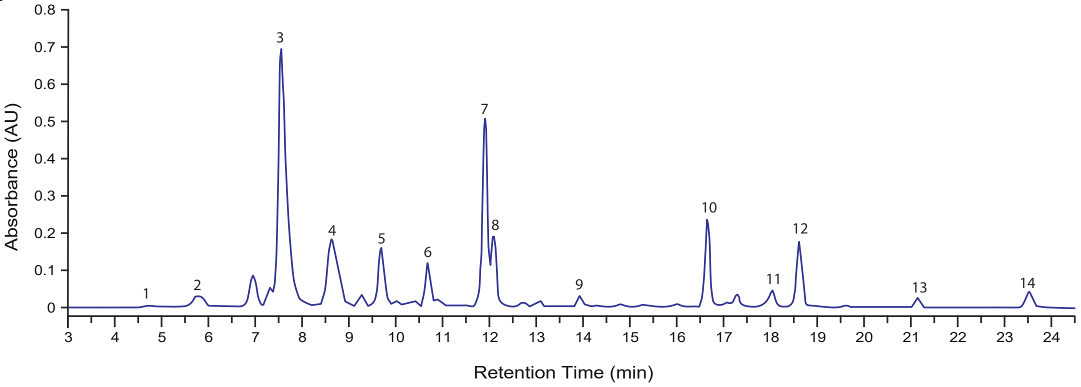

**b**

| Peak | Identification | Retention Time (min) | UV-Vis Absortion maxima (nm) |
|---|---|---|---|
| 1 | ND | 5.773 | 472 |
| 2 | ND | 6.547 | 422, 445, 470 |
| 3 | ND | 7.56 | 450, 477, 508 |
| 4 | Dinoxanthin | 8.627 | 418, 442, 471 |
| 5 | Diatoxanthin | 9.693 | 426, 449, 478 |
| 6 | ND | 10.687 | 431, 454, 483 |
| 7 | Lutein | 11.9 | 423, 448, 475 |
| 8 | Zeaxanthin | 12.097 | 428, 450, 478 |
| 9 | Canthaxanthin | 13.927 | 473 |
| 10 | Chlorophyll b | 16.647 | 458, 597, 645 |
| 11 | Chlorophyll a | 18.033 | 382, 412, 431, 617, 662 |
| 12 | ND | 18.613 | 461 |
| 13 | Pheophytin a | 21.14 | 409, 506, 537, 608, 665 |
| 14 | β-carotene | 23.51 | 431, 454, 481 |

**Extended Data Fig. 1. Characterization of Lake Kinneret chromophore extract. a**, HPLC profile of the chromophore extract. **b**, UV-Vis spectral characterization and identification of the peaks detected in (**a**).

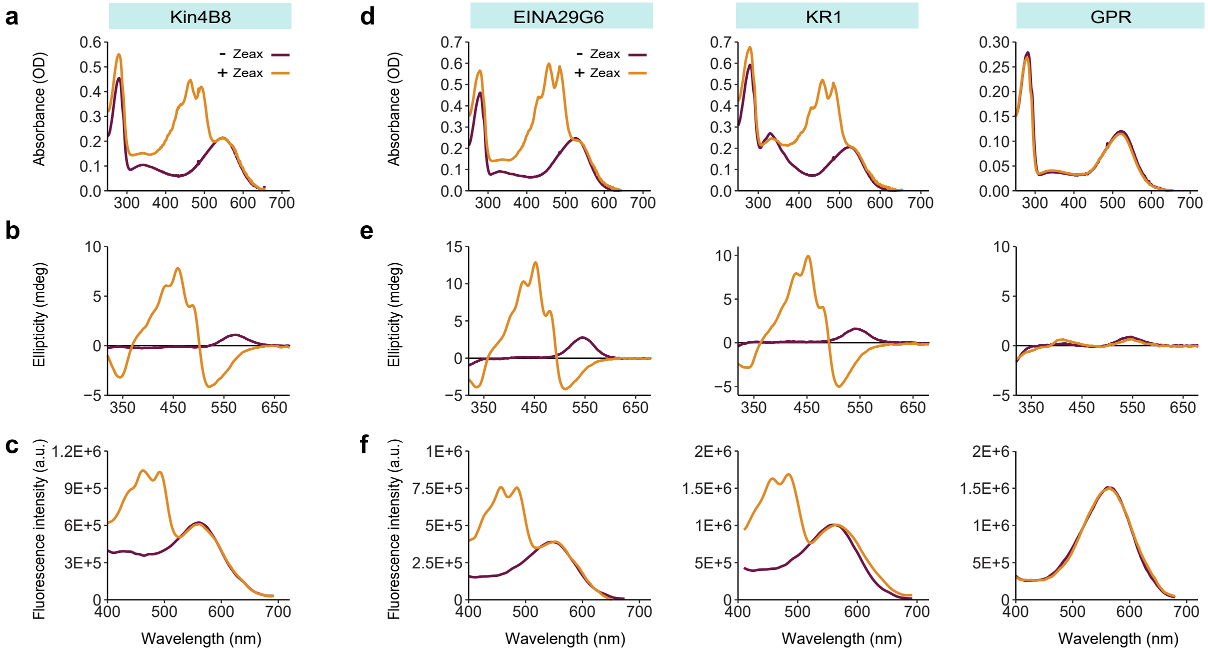

**Extended Data Fig. 2. Spectroscopic characterization of diverse rhodopsins bound to zeaxanthin. a** and **d**, Absorbance change of different rhodopsins upon incubation with zeaxanthin (Zeax). **b**, **d**, CD spectra with and without zeaxanthin. **c**, **f**, Fluorescence excitation spectra with and without zeaxanthin; emission monitored at 720 nm. The blue line represents the fluorescence excitation spectra of pure zeaxanthin; emission monitored at 720 nm.

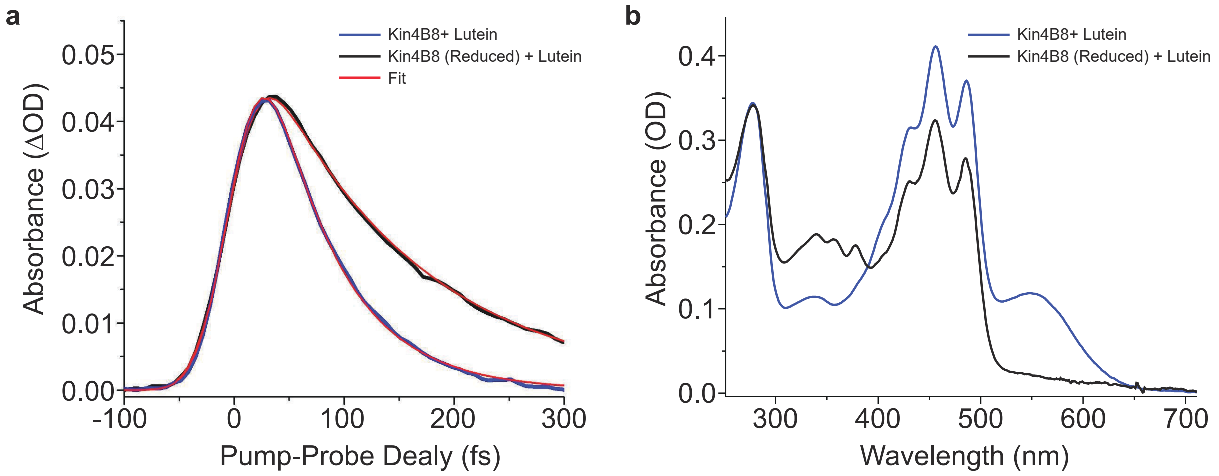

**Extended Data Fig. 3. Ultrafast spectroscopy characterization of lutein in complex with Kin4B8.** **a**, $S_2$ state's decay of lutein. Black- Kin4B8 (reduced RPSB bond)-lutein, Blue- Kin4B8-lutein. Red- their fit. X-axis presents the delay between pump and probe. Y-axis shows probe absorption difference in the presence and absence of pump pulse. Kinetic data were fitted with a function convolving 40 fs gaussian IRF and a mono-exponential decay. **b**, Reduction of retinal protonated Schiff-base by $NaBH_4$ in Kin4B8-lutein complex. Black-Absorption spectrum of Kin4B8-Lutein (before reduction). Red-Absorption spectrum of Kin4B8-Lutein (after reduction).

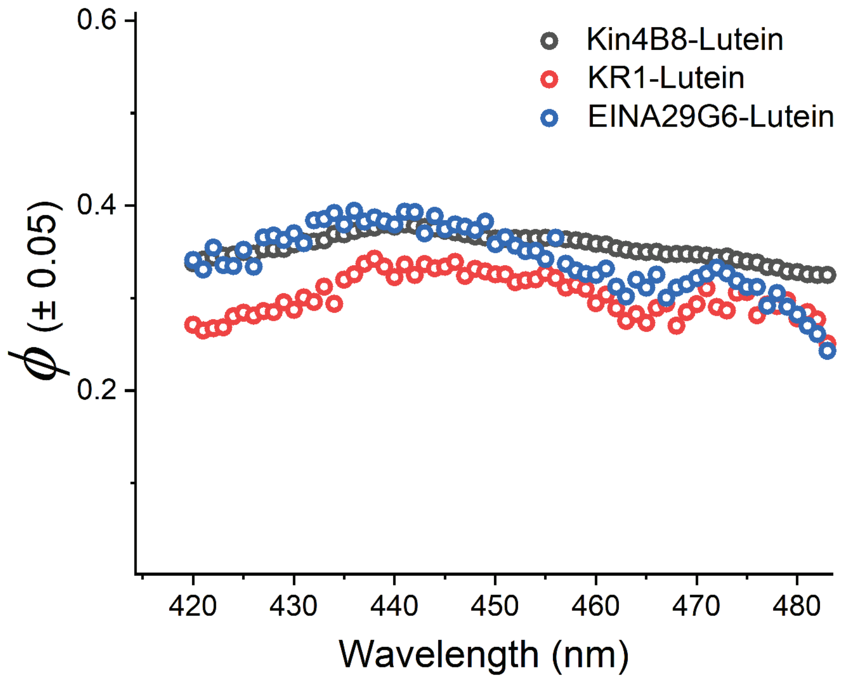

**Extended Data Fig. 4. Quantum efficiency of EET from lutein to different rhodopsins in complex, as a function of wavelength.**

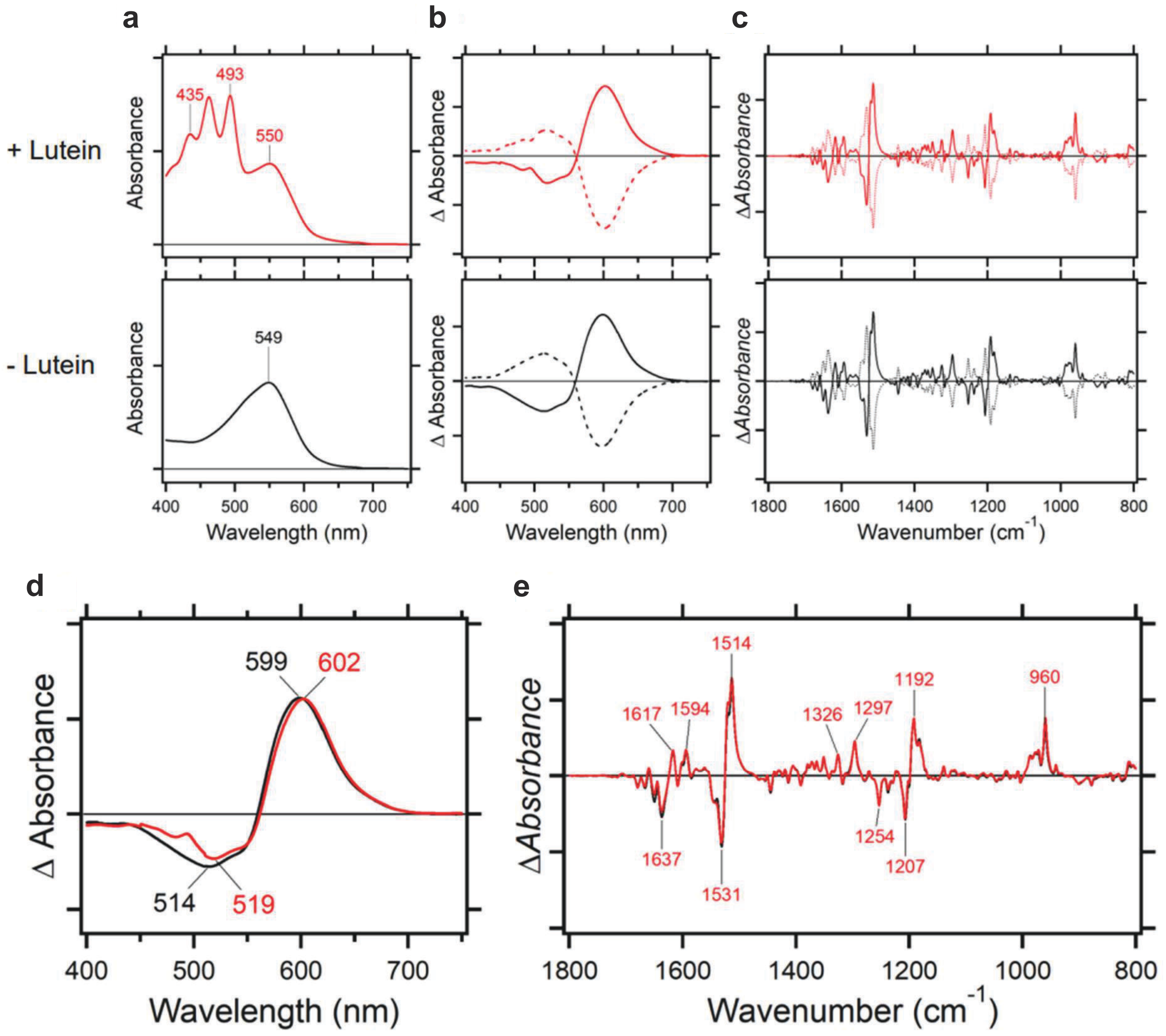

**Extended Data Fig. 5. FTIR Influence of lutein on the retinal photoisomerization in Kin4B8 at 77 K. a**, UV-visible absorption spectra of lipid-reconstituted Kin4B8 with (top) and without (bottom) lutein at 77 K. One division of the y-axis corresponds to 0.5 absorbance units. **b**, Difference UV-visible spectra upon illumination of Kin4B8 with (top) and without (bottom) lutein. Hydrated films of lipid-reconstituted Kin4B8 were first illuminated at 540 nm light (solid lines), followed by illumination at >590 nm light (broken lines) at 77 K. Solid and broken lines are mirror-imaged, indicating photochromic properties for Kin4B8 and the K intermediate. One division of the y-axis corresponds to 0.05 absorbance units. **c**, Difference FTIR spectra upon illumination of Kin4B8 with (top) and without (bottom) lutein. Hydrated films of lipid-reconstituted Kin4B8 with $H_2O$ were first illuminated at 540 nm light (solid lines), followed by illumination at >590 nm light (dotted lines) at 77 K. One division of the y-axis corresponds to 0.002 absorbance units. **d, e**, Light-induced difference UV-visible (**d**) and FTIR (**e**) spectra of Kin4B8 with (red) and without (black) lutein, where positive and negative signals originate from the K intermediate and unphotolyzed Kin4B8, respectively.

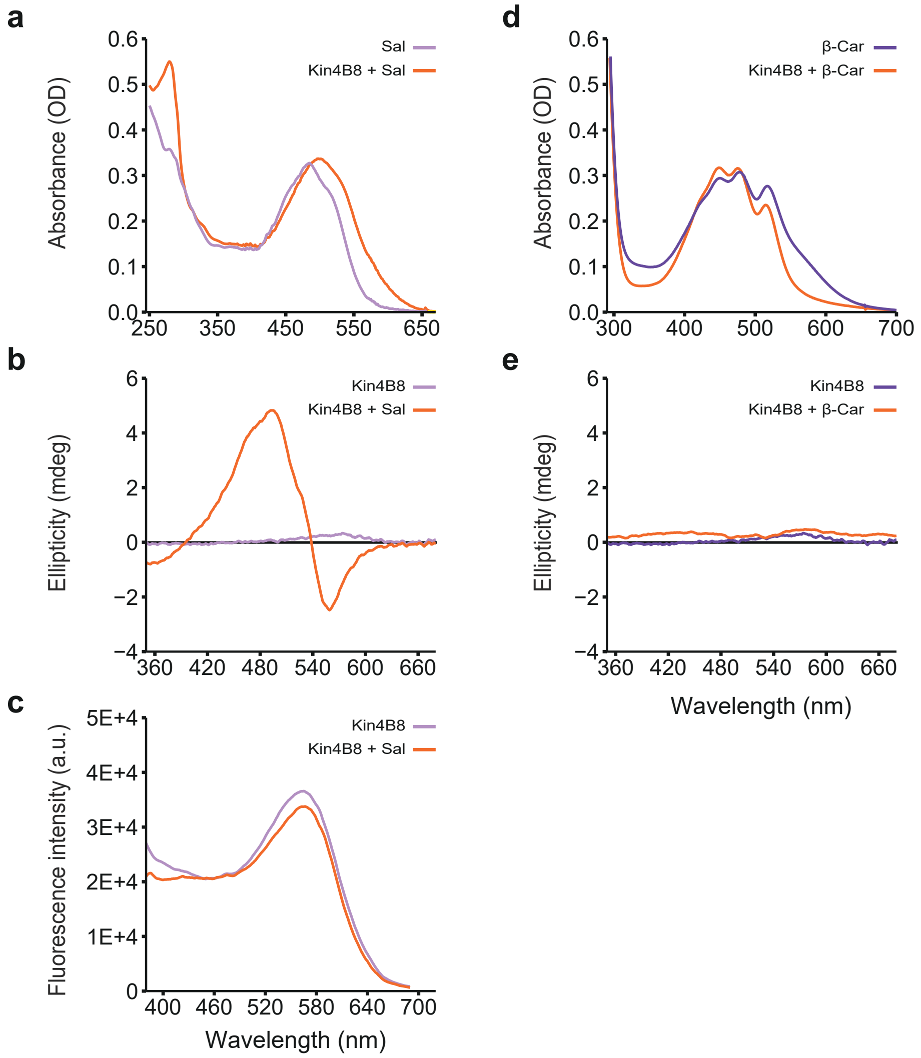

**Extended Data Fig. 6. Spectroscopic characterization of Kin4B8 bound to salinixanthin and β-carotene.** **a**, **d**, Absorption spectra of Kin4B8 with salinixanthin (Sal) or β-carotene (β-car), respectively. **b**, **e**, CD spectra with and without salinixanthin or β-carotene, respectively. **c**, Fluorescence excitation spectra with and without salinixanthin; emission monitored at 720 nm.

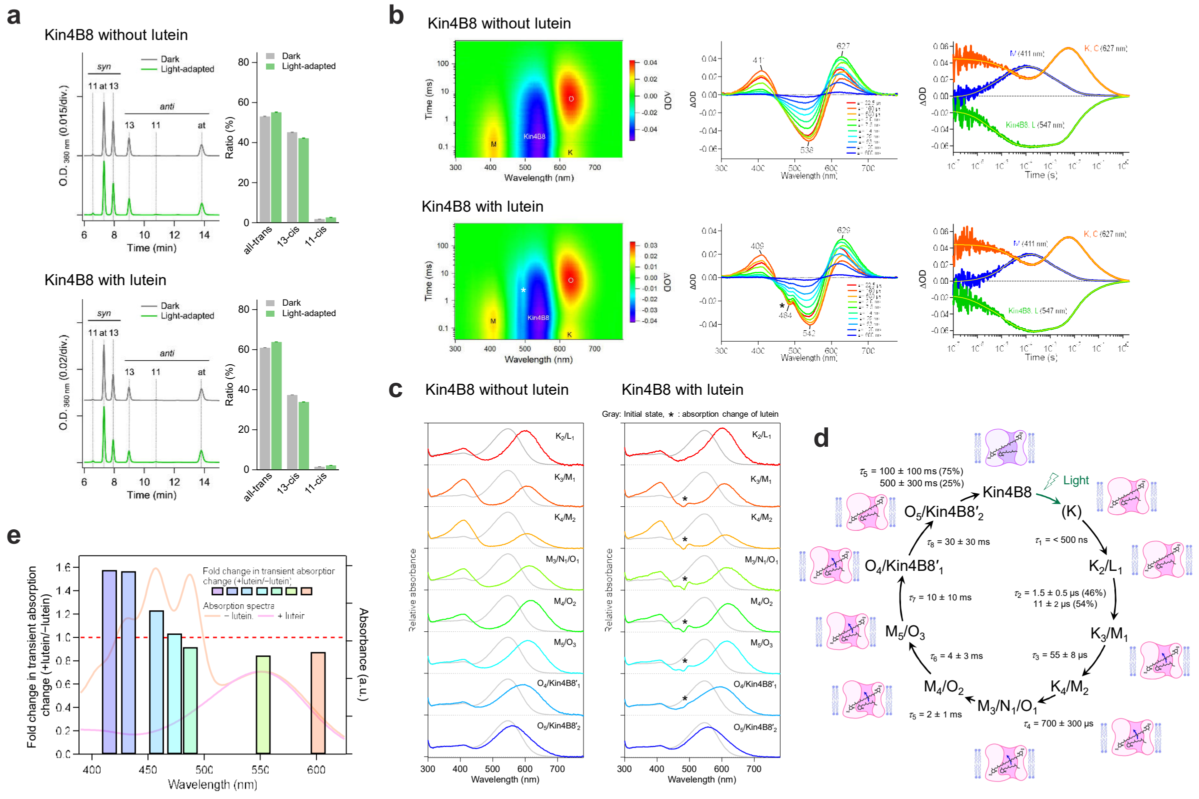

**Extended Data Fig. 7. The photocycle of Kin4B8.** **a**, Chromatogram of HPLC analyses (left) and the compositions of the retinal isomers (right) in Kin4B8 with (top) and without (bottom) lutein under the dark (gray) and light-adapted (green) conditions. at, 11, 13, syn, and anti indicate all-*trans*, 11-*cis*, 13-*cis*, *syn*, and *anti* configulations, respectively. **b**, Two-dimensional plot of transient absorption change (left), transient absorption spectra at different time points (middle), and time course of the transient absorption change (right) of Kin4B8 without (top) and with (bottom) lutein. Peaks derived from the absorption change of lutein are indicated by an asterisk. **c**, Absorption spectra of the photointermediates of Kin4B8 with (left) and without (right) lutein. Peaks derived from the absorption change of lutein are indicated by an asterisk. **d**, Photocycle model of Kin4B8. Conformational change of rhodopsin affects the structure of lutein (blue arrow) from $K_3/M_1$ to $O_4$/Kin4B8′$_2$. **e**, The ratios of transient absorption change in Kin4B8 with and without lutein at different excitation wavelengths (415, 432, 457, 473, 487, 552, and 601 nm) (bars colored according to the color of excitation light). The absorption spectra of Kin4B8 without (pink line) and with (orange line) lutein were overlaid. The red dashed line indicates no difference between with and without lutein.

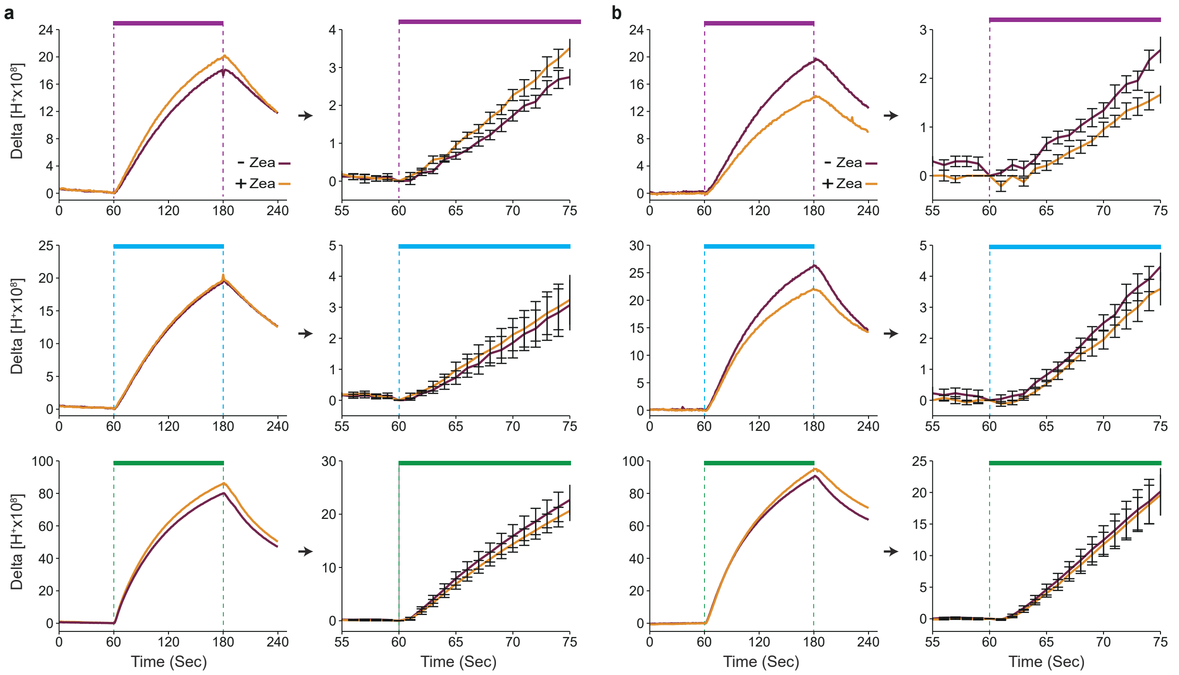

**Extended Data Fig. 8. Light-induced proton-pumping activity of Kin4B8 and Kin4B8-G153F. a** and **b**, Monitoring of pH changes in *E. coli* spheroplasts suspension expressing Kin4B8 or Kin4B8-G153F, respectively, with and without zeaxanthin. The spheroplasts were illuminated with violet (430 nm), blue (450 nm), or green (550 nm) light for 2 min (indicated by the colored bars). An enlarged plot of the first 15 sec of illumination is displayed to the right of each measurement. The presented traces are the average of six or more independent biological replicates.

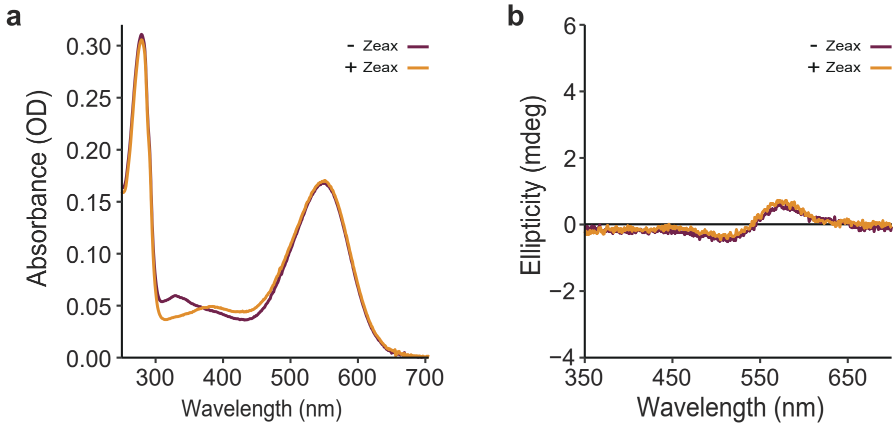

**Extended Data Fig. 9. Spectroscopic characterization of Kin4B8-G153F. a**, Absorption spectra with and without zeaxanthin. **b**, CD spectra with and without zeaxanthin**.**

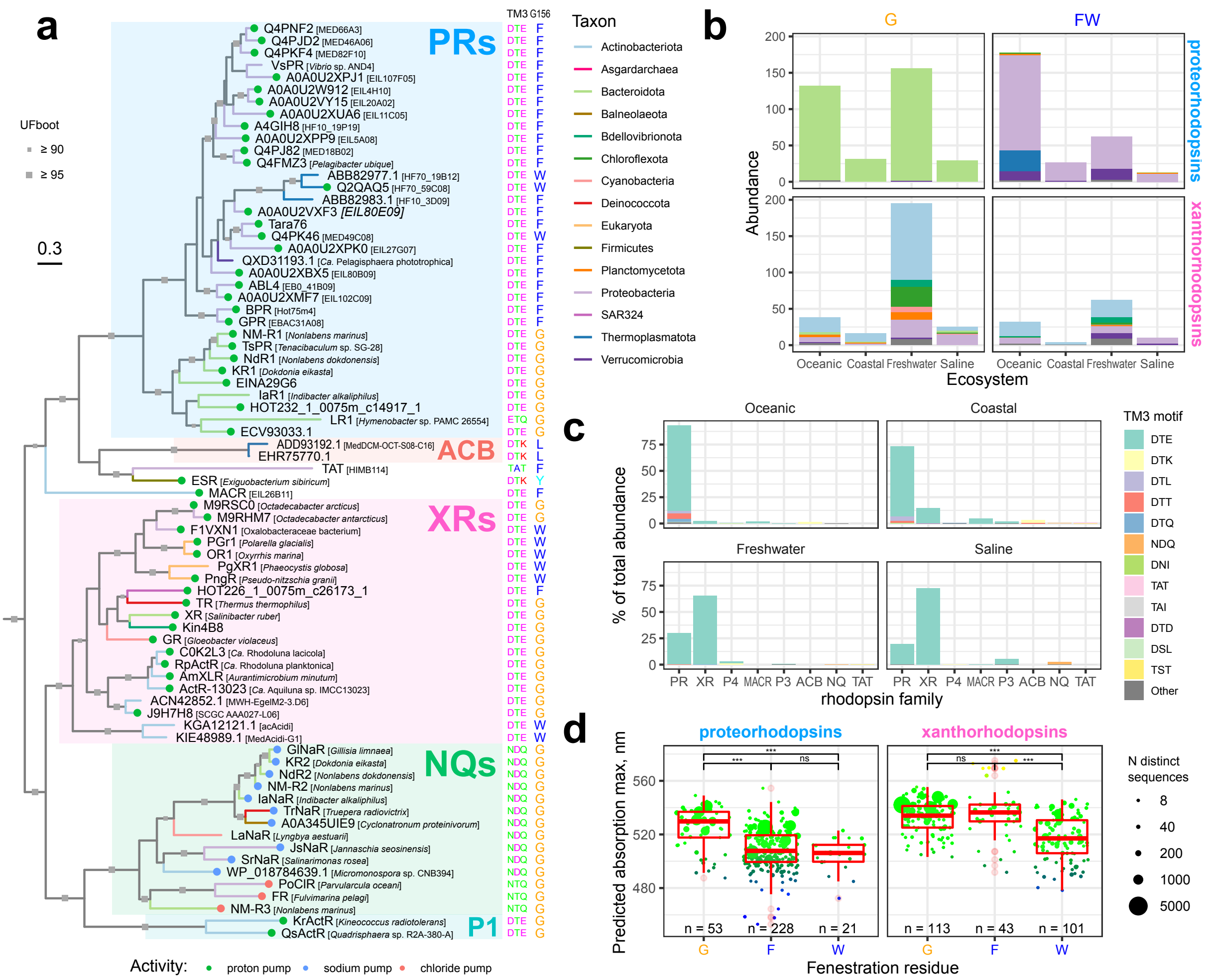

**Extended Data Fig. 10. Diversity and distribution of PRs and XRs with (G) and without (FW) fenestration among different prokaryotic phyla across four environments. a**, Phylogenetic analysis of the PR-XR clade based on representative protein sequences. Characterized ion pumps are indicated with dots, terminal branches are colored by the corresponding phylum. Major clades with more than one representative are highlighted and labeled. The tree is outgroup rooted. **b**, Distribution of PRs and XRs with the canonical TM3 motif DTE among genomes assigned to different taxa, with (G) and without (FW) fenestration. The analysis is based on GEM genomes and the numbers are summarized per operational taxonomic unit (OTU). The colors are as in panel (A). **c**, Relative abundance of different families of the clade across four habitats based on the metagenomic data in IMG/M. Only families with a total relative abundance of >0.1% are shown. **d**, Predicted absorption maxima for PRs and XRs with the three most frequent residues at the fenestration position. Individual observation corresponds to an average absorption maximum predicted with the rhodopsin BLASSO model for sequences with the same 24 residues of the retinal binding pocket[46]. The sequences from OM-RGC, IMG/M and GEM were pooled together. The size of the dots is proportional to the number of distinct rhodopsin domain sequences and the color approximates the predicted mean absorption spectra. Statistical differences between the groups were assessed with Dunn's test with FDR correction. Significance levels are indicated with asterisks: *** – adjusted $p$-values < 0.001. Abbreviation of family names in (A) and (C): ACB – Archaea clade B, ESR – *Exiguobacterium sibiricum* rhodopsin, NQ – NQ sodium and chloride pumps, MACR – marine actinobacteria clade rhodopsins, PR – proteorhodopsins, TAT – TAT rhodopsins, XR – xanthorhodopsins, P1 – unnamed clade including QsActR, KrActR and related rhodopsins, P3 and P4 – currently unnamed clades.

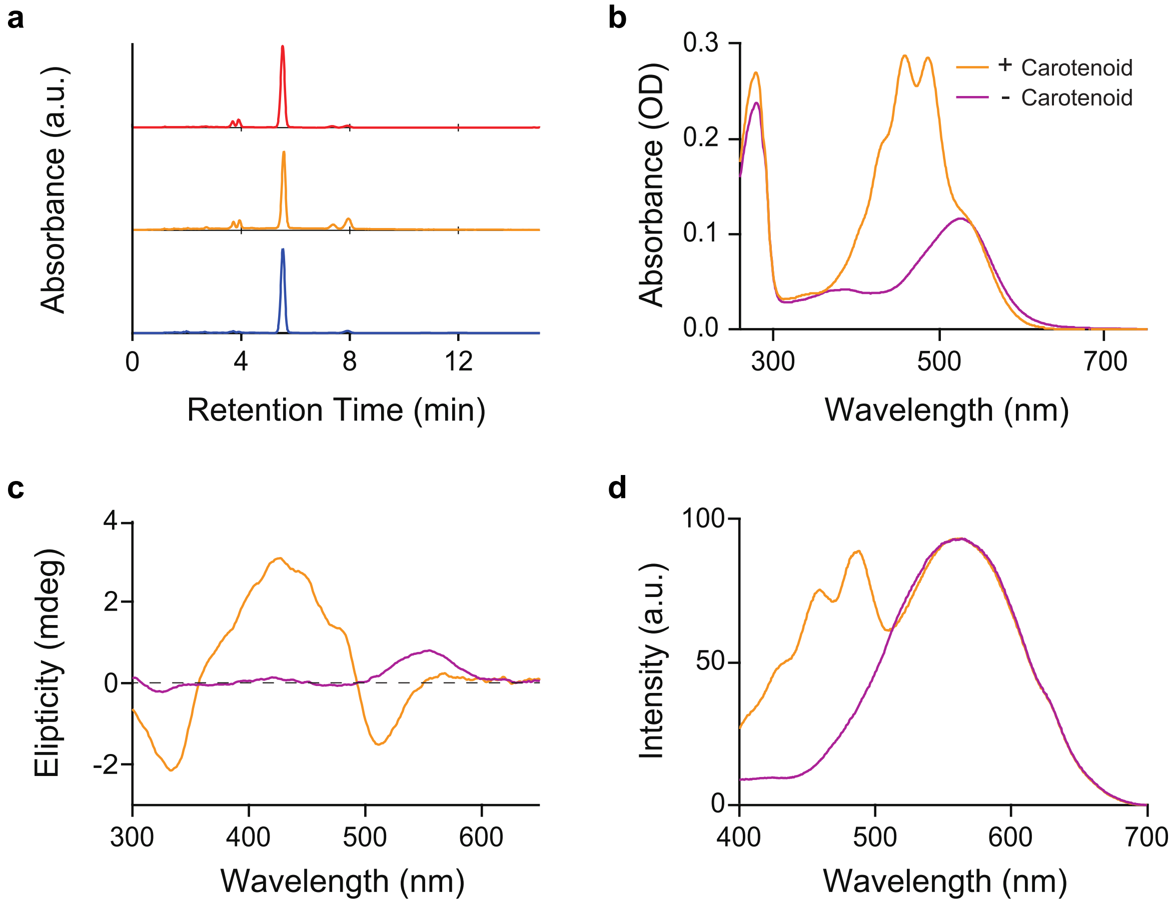

**Extended Data Fig. 11. Biophysical characterization of TsPR from *Tenacibaculum* sp. SG-28. a**, HPLC profiles of pigments, registered at 475 nm. Red, orange and blue chromatograms are whole carotenoid pigments extracted from SG-28 cells, binding carotenoid pigments to TsPR, and reference samples of pure zeaxanthin, respectively. **b**, Absorbance spectra of purified TsPR before (purple) and after (orange) adding whole carotenoid pigments extracted from SG-28 cells. **c**, CD spectra with and without zeaxanthin. **d**, Fluorescence excitation spectra with and without zeaxanthin; emission monitored at 720 nm.

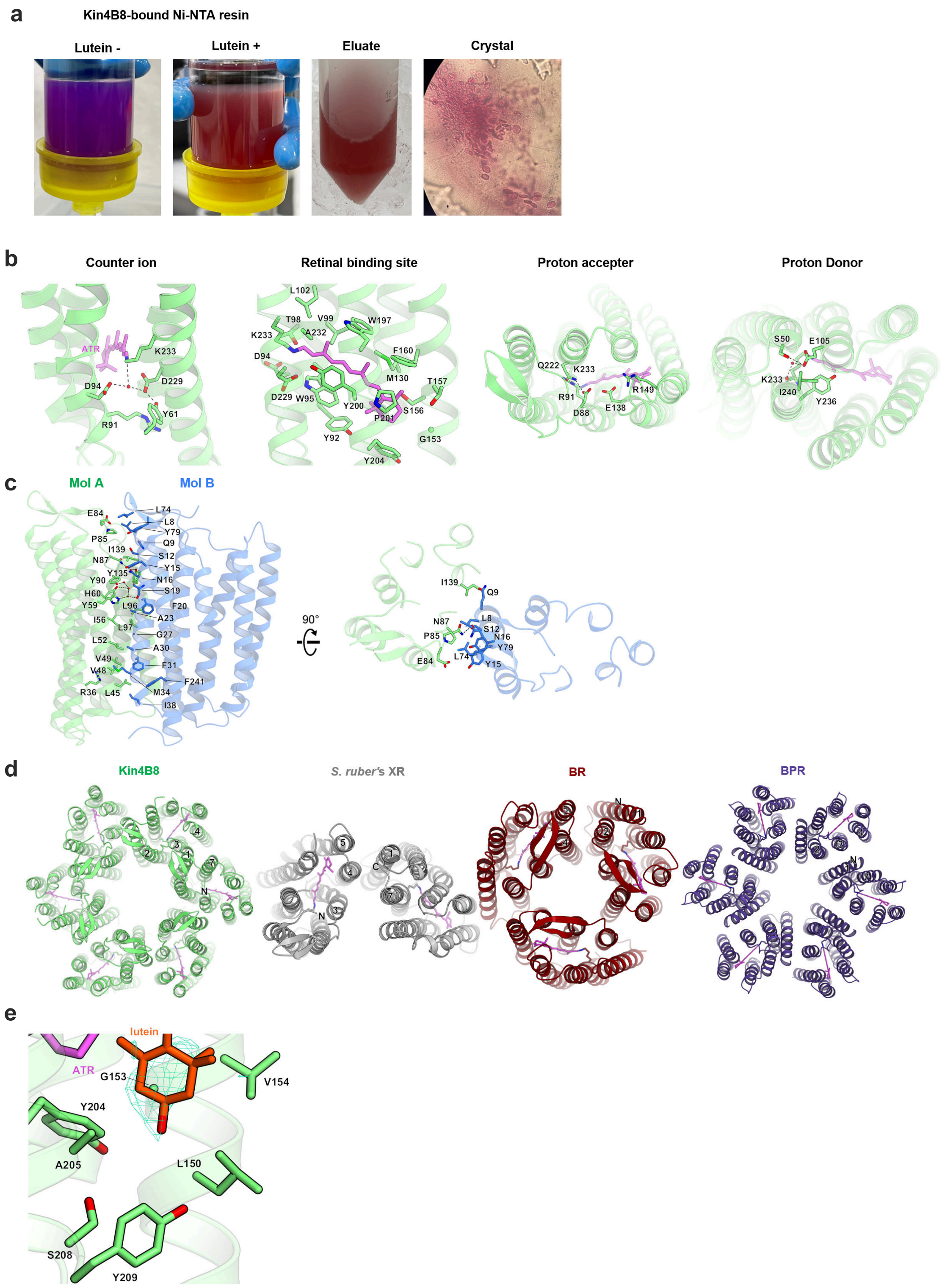

**Extended Data Fig. 12. Structural features of Kin4B8.** **a**, Kin4B8 color during purification and crystallization. We confirmed that Kin4B8 turns red upon lutein binding and keeps its color in the crystals. **b**, Key rhodopsin proton pump motifs in Kin4B8. **c**, Oligomeric interface between molA and molB. **d**, Comparison of the oligomeric structures of Kin4B8, *S. ruber*'s XR (PDB ID: 3DDL), BR (PDB ID: 1C3W), BPR (PDB ID: 4JQ6). With the ECL1 sheet inside, Kin4B8 forms a hexamer with aligned directions to the membrane in the crystal packing. The hexameric structure would reflect a physiological condition, in contrast to the previously reported head-to-tail dimer of *S. ruber*'s XR. **e**, Fo–Fc omit map for lutein contoured at 3.0σ around the fenestration.

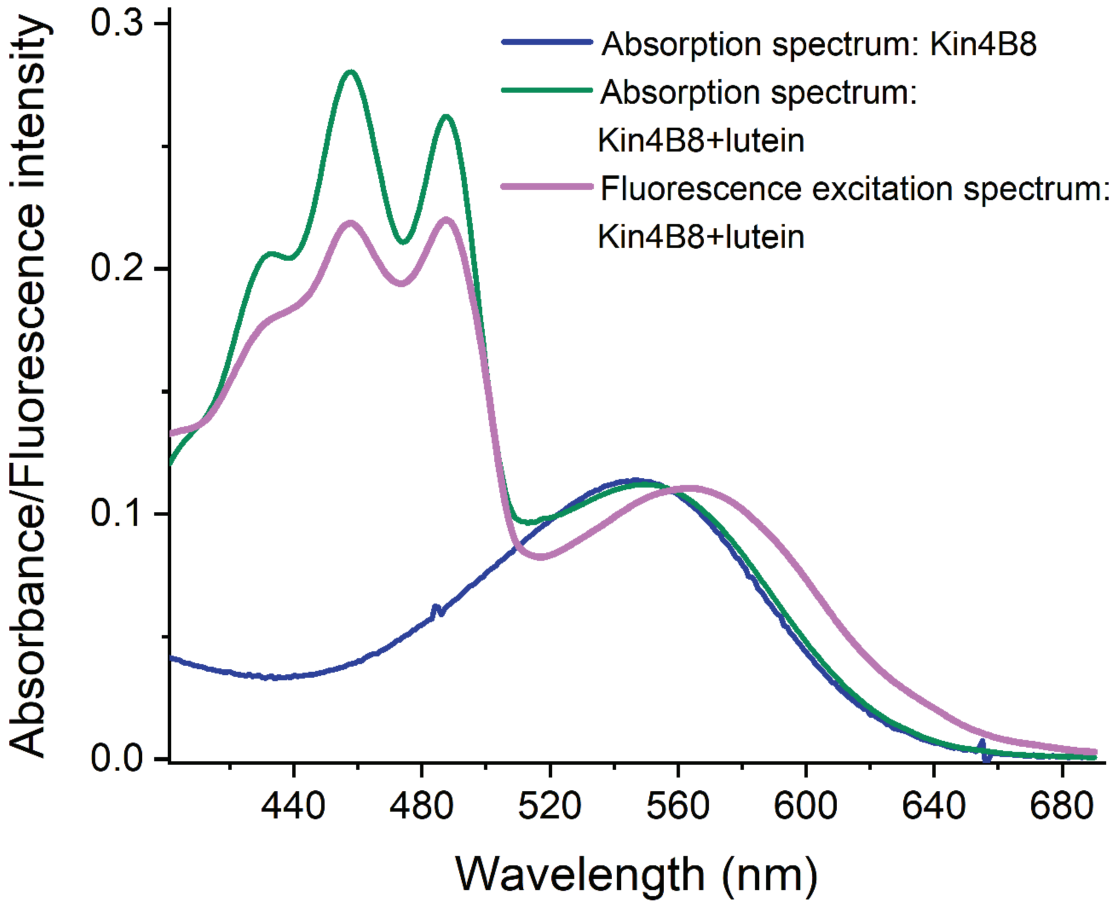

**Extended Data Fig. 13. Absorption and fluorescence excitation spectra of Kin4B8-lutein complex.** Fluorescence excitation spectrum scaled to the absorbance of the retinal component. Blue- absorption spectrum of retinal component, Green- absorption spectrum of Kin4B8-lutein complex, and Pink- fluorescence excitation spectrum of the Kin4B8-lutein complex.

**Extended Data Table 1. Quantum efficiency of EET from carotenoids to rhodopsins.**

| Rhodopsin | Carotene | % EET Efficiency (±5) |
|---|---|---|
| Kin4B8 | Lutein | 39 |
| | Zeaxanthin | 40 |
| KR1 | Lutein | 33 |
| | Zeaxanthin | 35 |
| EINA29G6 | Lutein | 42 |
| | Zeaxanthin | 40 |
| GPR | Lutein | 0 |
| | Zeaxanthin | 0 |

**Extended Data Table. 2. Data collection and refinement statistics.**

| | Kin4B8 |
|---|---|
| **Data collection** | |
| Space group | P321 |
| Cell dimensions | |
| *a, b, c* (Å) | 100.0, 100.0, 116.3 |
| α, β, γ (°) | 90, 90, 120 |
| Resolution (Å)* | 48.27-3.0 (3.107-3.0) |
| $R_{meas}$* | 0.5168 (8.544) |
| $<I/\sigma(I)>$* | 10.25 (0.93) |
| $CC_{1/2}$* | 0.994 (0.378) |
| Completeness (%)* | 99.51 (98.3) |
| Redundancy* | 23.0 (24.0) |
| **Refinement** | |
| Resolution (Å) | 43.88-3.0 |
| No. reflections | 13,912 |
| $R_{work}$ / $R_{free}$ | 0.2168 / 0.2547 |
| No. atoms | |
| Protein | 3898 |
| Lipid and chromophore | 225 |
| Water | 25 |
| Averaged *B*-factors (Å$^2$) | |
| Protein | 66.01 |
| Lipid and chromophore | 95.78 |
| Water | 53.56 |
| R.m.s. deviations from ideal | |
| Bond lengths (Å) | 0.003 |
| Bond angles (°) | 0.68 |
| Ramachandran plot | |
| Favored (%) | 97.2 |
| Allowed (%) | 2.6 |
| Outlier (%) | 0.2 |

*Values in parentheses are for the highest-resolution shell.
